# Selective Serotonin Reuptake Inhibitors Within Cells: Temporal Resolution in Cytoplasm, Endoplasmic Reticulum, and Membrane

**DOI:** 10.1101/2022.08.09.502705

**Authors:** Aaron L. Nichols, Zack Blumenfeld, Laura Luebbert, Hailey J. Knox, Anand K. Muthusamy, Jonathan S. Marvin, Charlene H. Kim, Stephen N. Grant, David P. Walton, Bruce N. Cohen, Rebekkah Hammar, Loren L. Looger, Per Artursson, Dennis A. Dougherty, Henry A. Lester

## Abstract

Selective serotonin reuptake inhibitors (SSRIs) are the most prescribed treatment for individuals experiencing major depressive disorder (MDD). The therapeutic mechanisms that take place before, during, or after SSRIs bind the serotonin transporter (SERT) are poorly understood, partially because no studies exist of the cellular and subcellular pharmacokinetic properties of SSRIs in living cells. We studied escitalopram and fluoxetine using new intensity-based drug-sensing fluorescent reporters (“iDrugSnFRs”) targeted to the plasma membrane (PM), cytoplasm, or endoplasmic reticulum (ER) of cultured neurons and mammalian cell lines. We also employed chemical detection of drug within cells and phospholipid membranes. The drugs attain equilibrium in neuronal cytoplasm and ER, at approximately the same concentration as the externally applied solution, with time constants of a few s (escitalopram) or 200-300 s (fluoxetine). Simultaneously, the drugs accumulate within lipid membranes by ≥ 18-fold (escitalopram) or 180-fold (fluoxetine), and possibly by much larger factors. Both drugs leave cytoplasm, lumen, and membranes just as quickly during washout. We synthesized membrane-impermeant quaternary amine derivatives of the two SSRIs. The quaternary derivatives are substantially excluded from membrane, cytoplasm, and ER for > 2.4 h. They inhibit SERT transport-associated currents 6- or 11-fold less potently than the SSRIs (escitalopram or fluoxetine derivative, respectively), providing useful probes for distinguishing compartmentalized SSRI effects. Although our measurements are orders of magnitude faster than the “therapeutic lag” of SSRIs, these data suggest that SSRI-SERT interactions within organelles or membranes may play roles during either the therapeutic effects or the “antidepressant discontinuation syndrome”.

**SIGNIFICANCE STATEMENT:** Selective serotonin reuptake inhibitors stabilize mood in several disorders. In general, these drugs bind to the serotonin (5-hydroxytryptamine) transporter (SERT), which clears serotonin from CNS and peripheral tissues. SERT ligands are effective and relatively safe; primary care practitioners often prescribe them. However, they have several side effects and require 2 to 6 weeks of continuous administration until they act effectively. How they work remains perplexing, contrasting with earlier assumptions that the therapeutic mechanism involves SERT inhibition followed by increased extracellular serotonin levels. This study establishes that two SERT ligands, fluoxetine and escitalopram, enter neurons within minutes, while simultaneously accumulating in many membranes. Such knowledge will motivate future research, hopefully revealing where and how SERT ligands “engage” their therapeutic target(s).

## INTRODUCTION

The approval of fluoxetine (Prozac®), the first serotonin reuptake inhibitor (SSRI), in 1986 transformed treatment of major depressive disorder (MDD). Prescribed regimens for MDD still use fluoxetine and other SSRIs, such as escitalopram (Lexapro®) (Wong et al., 1974; Wong et al., 1995; Beasley et al., 2000; Baldwin, 2002; Burke et al., 2002; Lepola et al., 2003; Lalit et al., 2004; Wong et al., 2005; Rao, 2007; Kennedy et al., 2009).

However, fascinating neuroscientific questions remain about the mechanism(s) of SSRI therapy. Answering the question, “Where do SSRIs go?” is necessary, but admittedly not sufficient, to address uncertainties surrounding SSRI mechanisms and modes of action. Therefore, this paper develops and exploits tools to study movements of fluoxetine and escitalopram within cells.

The experiments were conducted against the background of four non-mutually exclusive mechanisms that might explain SSRI action, presented in order of increasing novelty. First, the eponymous inhibition of the plasma membrane serotonin transporter (SERT) would increase extracellular serotonin levels, eventually causing (via unexplained intracellular mechanisms) amelioration of the clinical symptoms of MDD (Clevenger et al., 2018); we have termed this the “outside-in” mechanism (Lester et al., 2012). While serotonin levels in the synaptic cleft are rapidly altered after SSRI administration, a “therapeutic lag” of 2–6 weeks indicates a more complex mode of action (Nierenberg et al., 2000; Belmaker and Agam, 2008; Malhi and Mann, 2018). Depletion of serotonin in healthy individuals does not produce depressive effects, also suggesting that increased extracellular serotonin may be only one component of SSRI action (Salomon et al., 1997).

Second, the therapeutic lag may result mostly from a 10-day (or longer) duration of SSRI levels, eliminating the necessity for complex “outside-in” mechanisms but raising fundamental questions about SSRI pharmacokinetics. The steady-state cerebrospinal fluid (CSF) concentration in patients during escitalopram and fluoxetine treatment are 17–115 nM (Paulzen et al., 2016) and 13 ± 6 µM (Karson et al., 1992; Renshaw et al., 1992; Bolo et al., 2000) respectively (though the CSF fluoxetine concentration is confounded by simultaneous detection of both fluoxetine and its metabolite norfluoxetine). The apparent volume of distribution values for escitalopram (20 L kg^-1^) (Sogaard et al., 2005) and fluoxetine (20‒42 L kg^-1^) (Lee-Kelland et al., 2018) indicate substantial accumulation of each drug in tissues. Volume of distribution is sometimes correlated with increasingly positive logD_pH7.4_ values; escitalopram (1.41) and fluoxetine (1.83) follow this trend.

Third, vaguely defined “inside-out” mechanisms postulate that SSRIs enter the organelles of the early exocytotic pathway, where binding to nascent SERT may induce unconventional mechanisms such as pharmacological chaperoning (Lester et al., 2012). Thus, “inside-out” mechanisms would begin with SSRI-SERT binding, but in compartments distinct from the plasma membrane and via effects other than 5-HT reuptake. These mechanisms would cause the therapeutic lag.

Fourth, SSRIs may relieve MDD through additional mechanisms, involving targets other than SERT. Possible pathways include activation of the brain-derived neurotrophic factor (BDNF) receptor tyrosine kinase reporter 2 (TRKB) (Casarotto et al., 2021), and lipid rafts (Senese and Rasenick, 2021).

We used several methods. Analogous to recent work on the subcellular pharmacokinetics of other neuropsychiatric drugs (Bera et al., 2019; Shivange et al., 2019; Muthusamy et al., 2022; Nichols et al., 2022), we developed and applied new intensity-based drug sensing fluorescent reporters (“iDrugSnFRs”) for SSRIs, targeted to the plasma membrane (PM), cytoplasm, and endoplasmic reticulum (ER). We detected fluorescence changes resulting from ∼ 1 s solution changes, at cultured neurons and HeLa cells. We chemically detected drug accumulation by cultured cells and phospholipid-coated beads. We also synthesized membrane-impermeant quaternary amine derivatives of escitalopram and fluoxetine and compared their movements and SERT pharmacology to those of the SSRIs.

The subcellular pharmacokinetic data are more complex than expected from biophysical and biochemical studies on the iDrugSnFRs in solution or from our previous work on other central nervous system (CNS)-acting drugs. The data provide insight into three of the four potential mechanisms of SSRI action in the CNS.

## Materials and Methods

### Experimental design and statistical analysis

All dose response experiments using purified iDrugSnFR protein were performed in triplicate. The standard deviation was calculated for each data point acquired. Isothermal titration calorimetry (ITC) experiments were performed in triplicate and the standard error of the mean (SEM) was calculated. Half maximal inhibitory concentration (IC_50_) measurements for escitalopram, fluoxetine, and their quaternary derivatives were from a minimum of 10 cells, from which the SEM was calculated. Stopped-flow experiments were repeated 5 times and averaged, from which standard deviations were calculated (except for 100 s experiments, which were collected only once). Experiments in mammalian cell culture and mouse primary hippocampal culture were designed such that fluorescence response was averaged across a minimum of 5 cells from a minimum of two distinct fields of view, after which the SEM was calculated.

### Directed evolution of iDrugSnFR proteins using bacterial expression

Starting with iAChSnFR and intermediate biosensor constructs of that sensor, we constructed and optimized biosensors for each drug partner during iterative rounds of site-saturated mutagenesis (SSM) as previously described (Bera et al., 2019; Borden et al., 2019; Shivange et al., 2019; Unger et al., 2020). We utilized the 22-codon procedure including a mixture of three primers, creating 22 unique codons encoding the 20 canonical amino acids (Kille et al., 2013). The 22-codon procedure yields an estimated > 95% residue coverage for a collection of 96 randomly chosen clones.

A Spark M10 96-well fluorescence plate reader (Tecan, Männedorf, Switzerland) was used to measure resting and drug-induced fluorescence (F_0_ and ΔF, respectively). Bacterial lysates were tested with excitation at 485 nm and emission at 535 nm. Lysates were also measured against choline to evaluate potential endogenous intracellular binding. Promising clones were amplified and sequenced. The optimally responding construct in each round of SSM was used as a template for the next round of SSM.

S-slope allows for comparison between iDrugSnFRs with differing ΔF_max_/F_0_ values (Bera et al., 2019) at the beginning of the concentration-response relation, which is typically the pharmacologically relevant range. With lysates or purified protein, which allow complete concentration-response relations, the Hill coefficient is near 1.0. We therefore calculated

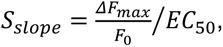

in units of µM^-1^.

### Measurements on purified iDrugSnFRs

Biosensors selected for further study were purified via the His_6_ sequence using an ÄKTA Start FPLC (GE Healthcare, Chicago, IL) as previously described (Shivange et al., 2019). Performance of protein quantification and concentration-response relations for drug-sensor partners was also as previously described (Shivange et al., 2019).

### Isothermal titration calorimetry (ITC)

ITC experiments were performed on an Affinity ITC (TA instruments, New Castle, DE) at 25 °C. Biosensor protein was buffer-exchanged into 3x PBS, pH 7.0. The SSRIs were dissolved in the same buffer. 450 µM escitalopram (Tocris, Bristol, United Kingdom) was titrated into 45 µM iEscSnFR and 700 µM N-N-dimethylfluoxetine was titrated into 140 µM iFluoxSnFR. Analysis, including correction for changes in enthalpy generated from the dilution of the ligands into the cell during titration, was performed using a single-site binding model in the manufacturer’s Nanoanalyze software.

### Stopped-flow kinetic analysis

Kinetics were determined by mixing equal volumes of 0.2 µM iDrugSnFR protein (in 3x PBS, pH 7.0) with varying concentrations of cognate ligand in an Applied Photophysics (Surrey, United Kingdom) SX20 stopped-flow fluorimeter with 490 nm LED excitation and 510 nm long-pass filter at room temperature (22 °C). “Mixing shots” were repeated 5 times and averaged (except for 100 s experiments, which were collected only once). Standard deviations are not included on the plots but are nearly the same size as the data markers. The first 3 ms of data were ignored because of mixing artifacts and account for the dead time of the instrument. Data were plotted and time courses were fitted, when possible, to a single exponential, with rate constant k_obs_. When the time course did not fit well to a single rising exponential, it was fitted to the sum of two increasing exponentials.

### Synthesis of N-methylescitalopram and N-N-dimethylfluoxetine

Synthesis of N-methylescitalopram was as previously published, with escitalopram replacing citalopram as the starting reagent (Bismuth-Evenzal et al., 2010). To generate N-N- dimethylfluoxetine, fluoxetine hydrochloride (Sigma Aldrich, St. Louis, MO) (60 mg, 0.174 mmol) was dissolved in MeCN (5 mL). Et_3_N (130 μL, 5 equiv.) was added, followed by MeI (324 μL, 30 equiv.). The reaction was stirred at room temperature for 20 min. EtOAC (10 mL) was added, and the resulting precipitate was removed by filtration. The filtrate was concentrated, dissolved in dichloromethane, and washed with water (3x). The organic layer was dried over MgSO_4_ and concentrated to give a yellow oil (44 mg, 54%). ^1^H-NMR (400 MHz, CDCl_3_): δ 7.46-7.29 (m, 7H), 6.97 (d, *J* = 9.05 Hz, 2H), 5.57 (dd, *J* = 6.32 Hz 1H), 4.22-4.15 (m, 1H), 3.80-3.73 (m, 1H), 3.38 (s, 9H), 2.41-2.35 (m, 2H). ESI: M^+^ calculated for C_19_H_23_F_3_NO^+^ 338.17, found 338.22.

### Expression in HeLa cells

We constructed three variants of each iDrugSnFR for expression in mammalian cells. The plasma membrane (suffix “_PM”) and endoplasmic reticulum (suffix “_ER”) variants were constructed by circular polymerase extension cloning (Quan and Tian, 2009). To create the _PM constructs, we cloned the bacterial constructs into pCMV(MinDis), a variant of pDisplay (ThermoFisher Scientific, Waltham, MA) lacking the hemagglutinin tag (Marvin et al., 2013). To generate the _ER constructs, we replaced the 14 C-terminal amino acids (QVDEQKLISEEDLN, including the Myc tag) with an ER retention motif, QTAEKDEL (Shivange et al., 2019). To generate the cytoplasm-targeted (suffix “_cyto”) variants, we used Gibson assembly (Gibson et al., 2009) to remove existing N- and C-terminal tags and incorporated an N-terminal strong nuclear exclusion sequence (NES) (DIDELALKFAGLDL) (Guttler et al., 2010).

We transfected the iDrugSnFR cDNA constructs into HeLa cells. Cell lines were purchased from ATCC (Manassas, VA) and cultured according to ATCC protocols. For chemical transfection, we utilized either Lipofectamine 2000 or Lipofectamine 3000 (ThermoFisher Scientific), following the manufacturer’s recommended protocol. Cells were incubated in the transfection medium for 24 h and imaged 24–48 h after transfection.

### AAV production and transduction in primary mouse hippocampal neuronal culture

The adeno-associated virus plasmid vector AAV9-hSyn was described previously (Challis et al., 2019). PM- and ER-targeted virus was purified using the AAVpro Purification Kit (TakaraBio USA). Cytoplasm-targeted virus was purified according to (Challis et al., 2019). Mouse embryo dissection and culture were previously described (Shivange et al., 2019). About 4 days after dissection, _ER constructs were transduced at an MOI of 0.5 to 5 x 10^4^, _PM constructs at an MOI of 0.5 to 1 x 10^5^, and _cyto constructs at an MOI of 5 x 10^4^. Neurons were imaged ∼2–3 weeks post-transduction.

### Time-resolved fluorescence measurements in live mammalian cells and primary mouse hippocampal neuronal culture

Time-resolved concentration-response imaging was performed on a modified Olympus IX-81 microscope (Olympus microscopes, Tokyo, Japan) in widefield epifluorescence mode using a 40X lens. Images were acquired at 2–4 frames/s with a back-illuminated EMCCD camera (iXon DU-897, Andor Technology USA, South Windsor, CT) controlled by Andor IQ3 software. Fluorescence measurements at λ_ex_ = 470 nm and the epifluorescence cube were as previously described (Srinivasan et al., 2011; Shivange et al., 2019).

Solutions were delivered from elevated reservoirs by gravity flow, via solenoid valves (Automate Scientific, Berkeley, CA), through tubing fed into a manifold, at a rate of 1–2 mL/min. The vehicle was HBSS. Other details have been described (Srinivasan et al., 2011; Shivange et al., 2019). Data analysis procedures included subtraction of “blank” (extracellular) areas and corrections for baseline drifts using Origin Pro 2018.

For folimycin incubation experiments, primary hippocampal culture dishes were incubated with 80 nM folimycin (Sigma Aldrich) for 10 min prior to the standard time-resolved concentration-response imaging outlined above (Tischbirek et al., 2012).

### Spinning disk confocal fluorescence images

HeLa cells and mouse primary hippocampal culture were transfected or transduced as described above. Live-cell images were collected using a Nikon (Melville, NY) Ti-2E spinning disk laser scanning confocal inverted microscope through a 100X objective, 1.49 NA (oil), 120 μm WD. The laser wavelength was 488 nm at 15% power. Dishes were imaged in a custom incubator (Okolab, Ottaviano, Italy) at 37° C and 5% CO_2_. Initial images were taken in HBSS. To add drug, we doubled the bath volume by adding HBSS containing drug, using a hand-held pipette. The final drug concentrations for each set of experiments were: 10 µM, escitalopram; and 10 µM, fluoxetine.

### Probing inhibition of hSERT activity using electrophysiology

Human serotonin transporter (hSERT) cDNA was transferred to the pOTV vector (a gift from Dr. Mark Sonders, Columbia University). The K490T mutation (Cao et al., 1997) was made using the QuikChange protocol (Agilent, Santa Clara, CA). cDNA was linearized with NotI digestion (New England Biolabs, Ipswitch, MA) and purified using the QiaQuick PCR Purification kit (Qiagen, Hilden, Germany). Purified DNA was then transcribed *in vitro* using the T7 mMessage Machine kit (Ambion, Austin, TX). *Xenopus laevis* oocytes were isolated, injected with cRNA (20 ng in 50 nL nuclease-free water), and incubated at 19 °C for 3 days in Ca^2+^-free ND96 solution (96 mM NaCl, 2 mM KCl, 1 mM MgCl_2_, 5 mM HEPES, pH 7.5) supplemented with 0.05 mg/mL gentamycin (Sigma Aldrich), 2.5 mM sodium pyruvate (Acros Organics, Geel, Belgium), and 0.67 mM theophylline (Sigma Aldrich).

Two-electrode voltage clamp electrophysiology was performed on an OpusXpress 6000A (Molecular Devices Axon Instruments, San Jose, CA) at 20–25 °C. All compound solutions were prepared using Ca^2+^-free ND96 solution (pH 5.5). Solution containing only SSRI or quaternary derivative (0.67 mL) was applied to oocytes over 10 s, followed by a 10 s incubation. Solution containing the same concentration of SSRI/quaternary derivative and 3 μΜ 5-HT (1 mL) was then applied over 15 s. This process was followed by a 3.2 min washout period at a buffer flow rate of 3 mL/min (Fig. 9 below).

Oocytes were impaled with borosilicate glass electrodes filled with 3 M KCl (0.3–3.0 MΩ resistance) and held at -60 mV. Ca^2+^-free ND96 solution (pH 7.5) was used as a running buffer. In Clampfit 10.3 (Molecular Devices Axon Instruments), we employed a low-pass Gaussian filter at 5 Hz, then subtracted the average baseline current preceding application of compound solutions in low-pH buffer. For each cell, peak currents at each dose were normalized to the maximum transport-associated current (Mager et al., 1994) measured by applying 3 μM 5-HT in the absence of inhibitor (Fig. 9 below). Normalized currents were then averaged and fitted to the Hill equation using Prism 9 (GraphPad Software, Inc., San Diego, CA).

### LogD calculations

We used Chemicalize (https://chemaxon.com/products/chemicalize) to calculate LogP and pK_a_. The software then calculates

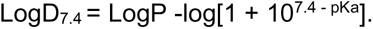

### Simulations of diffusion and binding

The simulation approximates Fick’s law as a sequence of fluxes between nested intracellular and extracellular shell compartments, governed by first-order rate constants k_f,_ = k_b_ (Yu et al., 2016). Each shell is treated as a well-stirred compartment. The units and dimensions are those used by Yu et al. (2016), including μm, fL (μm^3^), ms, and molarity (fM, μM, or M). Most shells have thickness 0.495 or 0.5, or 0.505 μm. For fluxes between shells, the effective diffusion constant is the free-solution value.

In classical analyses, for instance Yu et. al, 2016, the “membrane barrier” is not a shell; it is infinitely thin and has zero volume. In the model, the “membrane barrier” is located at a radius of 7.505 μm, and its permeability is represented by a single equal pair of rate constants k_f,_ = k_b_ for flux between one pair of neighboring shells.

The permeability of the “membrane barrier” is calculated as though it were a shell of finite volume: thickness 0.01 μm (10 nm), twice the value assumed by Kapoor et al., (2019) to account for proteins. The membrane permeability of the “membrane barrier” is calculated as though it were governed by free radial diffusion, as reduced by two large multiplicative factors. The first factor, n_pH_, accounts for the reduced availability of the neutral form of fluoxetine, given the difference between the pKa of fluoxetine and that of the cell. The second factor, n_accum_, is the reduction caused by binding to membrane lipids (Crank, 1975). The factor n_accum_ = (lipid molarity in the shell)/(assumed fluoxetine-lipid K_d_). The latter K_d_ is the most important adjustable parameter (see Results, Fig. 5C, Fig. 6L, and Table 1).

**Table 1.** Values from an Excel worksheet that calculates the parameters of the diffusion- binding model for neurons (see Methods). Another worksheet in the workbook calculates the model for HeLa cells. Several columns with intermediate calculations are hidden. The workbook, an .xlsx file, is at https://github.com/lesterha/lesterlab_caltech. The gray-shaded columns are the volumes of each shell and the bidirectional rate constants k_f,_ = k_b_ for the flux between each shell and the next larger shell (see Methods). The yellow row represents the calculations contributing to both the permeability of the “membrane barrier” and fluoxetine accumulation in the “membrane shell”. In the “membrane barrier”, the diffusion constant is reduced by two multiplicative factors (see Methods). The factor, n_pH_, accounts for the reduced availability of the neutral form of fluoxetine, given the difference between the calculated pKa of fluoxetine (9.8) and that of the external solution (7.4). The factor, n_accum_ is the (lipid molarity in the shell)/(assumed fluoxetine-lipid K_d_). The lipid molarity is calculated from the usual assumption that each membrane leaflet has a lipid density of 2.5 million molecules / μm^2^ (Alberts et al., 2015). The assumed fluoxetine-lipid K_d_ is the most important adjustable parameter. The value of 2.2 mM produces a half-time of 251 s and is consistent with the measured value of at ≥ 100 μM (Treyer et al., 2019). In the worksheet for HeLa, K_d_.has the value of 22 mM. The “membrane barrier” comprises a set of two equal rate constants, as though it were physically located at 7.505 μm. The blue-background row gives the rate constants corresponding to the permeability of the “membrane barrier”. The value of n_accum_ in the yellow row is also used to calculate accumulation in the “membrane shell”. Varying the assumed “membrane shell” thickness over a 3-fold range changed the simulated kinetics by < 10%, because the model’s structure has compensatory changes in several parameters.

|  |  |  | pH divisor | binding divisor | 1st-order rate constants | permeability |  |  |
| --- | --- | --- | --- | --- | --- | --- | --- | --- |
| compartment | shell outer radius | shell volume | n<sub>pH</sub> | n<sub>accum</sub> | k<sub>f</sub> =k<sub>b</sub> =<br>D<sub>eff</sub>*A / thickness | Classical permeability,<br>k = D<sub>eff</sub> / thickness | lipid molarity in shell | assumed fluoxetine-lipid K<sub>d</sub> |
|  | μm | μm<sup>3</sup> |  |  | μm<sup>3</sup>/ms | μm/ms | M | M |
| cytoplasmic shells | 0.5 | 0.52 | 1 | 1 | 1.26 | 0.4 |  |  |
|  | 1 | 3.67 | 1 | 1 | 5.03 | 0.4 |  |  |
|  | 1.5 | 9.95 | 1 | 1 | 11.31 | 0.4 |  |  |
|  | 2 | 19.37 | 1 | 1 | 20.11 | 0.4 |  |  |
|  | 2.5 | 31.94 | 1 | 1 | 31.42 | 0.4 |  |  |
|  | 3 | 47.65 | 1 | 1 | 45.24 | 0.4 |  |  |
|  | 3.5 | 66.50 | 1 | 1 | 61.58 | 0.4 |  |  |
|  | 4 | 88.49 | 1 | 1 | 80.42 | 0.4 |  |  |
|  | 4.5 | 113.62 | 1 | 1 | 101.79 | 0.4 |  |  |
|  | 5 | 141.90 | 1 | 1 | 125.66 | 0.4 |  |  |
|  | 5.5 | 173.31 | 1 | 1 | 152.00 | 0.4 |  |  |
|  | 6 | 207.87 | 1 | 1 | 181.00 | 0.4 |  |  |
|  | 6.5 | 245.57 | 1 | 1 | 212.37 | 0.4 |  |  |
|  | 7 | 286.41 | 1 | 1 | 246.30 | 0.4 |  |  |
|  | 7.495 | 326.86 | 1 | 1 | 282.37 | 0.4 |  |  |
| membrane shell | 7.505 | 7.07 | 230 | 208 | 0.01 | 8.36E-06 | 0.416 | 2.00E-03 |
| mebrane barrier |  | NA |  |  | 0.01 | 8.36E-06 |  |  |
| extracellular shells | 8 | 373.98 | 1 | 1 | 324.95 | 0.4 |  |  |
|  | 8.5 | 427.78 | 1 | 1 | 363.17 | 0.4 |  |  |
|  | 9 | 481.19 | 1 | 1 | 407.15 | 0.4 |  |  |
|  | 9.5 | 537.74 | 1 | 1 | 453.65 | 0.4 |  |  |
|  | 10 | 597.43 | 1 | 1 | 502.65 | 0.4 |  |  |
|  | 10.5 | 660.26 | 1 | 1 | 554.18 | 0.4 |  |  |
|  | 11 | 726.23 | 1 | 1 | 608.21 | 0.4 |  |  |
|  | 11.5 | 795.35 | 1 | 1 | 664.76 | 0.4 |  |  |
|  | 12 | 867.60 | 1 | 1 | 723.82 | 0.4 |  |  |
|  | 12.5 | 943.00 | 1 | 1 | 785.40 | 0.4 |  |  |
|  | 13 | 1021.54 | 1 | 1 | 849.49 | 0.4 |  |  |
|  | 13.5 | 1103.22 | 1 | 1 | 916.09 | 0.4 |  |  |
|  | 14 | 1188.04 | 1 | 1 | 985.20 | 0.4 |  |  |
|  | 14.5 | 1276.01 | 1 | 1 | 1056.83 | 0.4 |  |  |
|  | 15 | 1367.12 | 1 | 1 | 1130.97 | 0.4 |  |  |
|  | 15.5 | 1461.36 | 1 | 1 | 1207.63 | 0.4 |  |  |

Most classical analyses do not consider drug accumulation within the infinitely thin “membrane barrier” of zero volume. Therefore, we enhanced the classical simulation by including a routine that simultaneously calculates drug accumulation within a “membrane shell” of finite thickness and volume. The composition, thickness, composition, and volume are exactly those used to compute the permeability of the “membrane barrier”, above. Thus, the “membrane shell” has the lipid molarity and fluoxetine-lipid K_d_ described above and has a thickness of 10 nm (inner and outer radii at 7.495 and 7.505 μm respectively). Fluoxetine accumulation is calculated by simply multiplying the [fluoxetine] within the “membrane shell” by n_accum_. Fluoxetine accumulation does not deplete the total number of drug molecules. The inward-facing border of the “membrane shell” undergoes free diffusion with the next inward shell, as described for all other pairs of adjoining shells. The outer border of the “membrane shell” is the “membrane barrier”, whose permeability is described above. In the simulated results, the simulated waveform of drug concentration within the “membrane shell” is simultaneous with those in the cytoplasm, within ∼ 50 ms.

This conceptual scheme is valid only if the concentration source and sink lie outside the “membrane barrier”, allowing the accumulation within the “membrane shell” to be influenced by the delayed permeation through the “membrane barrier”. A more complete version, also allowing sources and sinks within the cell, would include both a 5 nm thick “inner membrane shell” and a 5 nm thick “outer membrane shell”, flanking the “membrane barrier”.

The model was constructed in the graphical user interface (GUI) of MATLAB Simbiology (Mathworks, Natick, MA). For our purposes, this interface has heuristic value; but it has the disadvantage that rate constants and shell volumes must be calculated externally. Therefore, we transferred the parameters manually to the GUI from the calculations and assumptions in an Excel spreadsheet (Table 1). Simbiology then integrated the equations to produce drug molarity vs time in each spherical shell (Fig. 5C and 6L). Both the Simbiology project (.sbproj) and the Excel spreadsheet may be downloaded from https://github.com/lesterha/lesterlab_caltech

For our purposes, Simbiology itself has the strengths (a) that it verifies consistency among the dimensions and units and (b) that it has robust routines for integrating stiff differential equations. Simbiology has the limitations (a) that it cannot treat surface densities in a compartment of zero volume and (b) that its dosing routines cannot jump the concentration of a source or sink. Therefore, the wash-in and washout phases of Fig. 5C and 6L were simulated separately.

### Total cellular accumulation, intracellular bioavailability, and lipid binding

Atorvastatin calcium salt, escitalopram oxalate, fluoxetine hydrochloride, lopinavir, and warfarin were obtained from Sigma-Aldrich at their highest degree of purity (≥ 98%). Atorvastatin and lopinavir were selected as reference compounds (Mateus et al., 2017) and warfarin was used as an internal standard. Atorvastatin, escitalopram, fluoxetine, lopinavir, and warfarin were made as stocks in DMSO (≥ 2 mM).

Total accumulation ratio (Kp) was measured as described previously (Treyer et al., 2019), but at several time points. In Dulbecco’s modified Eagle medium (DMEM) with addition of L-glutamine and 10% FBS, human embryonic kidney 293 (HEK293) cells were seeded at passage 14 at 6 x 10^5^ cells/mL in 24-well Corning Cellbind plates (Corning, NY). At confluence, they were washed twice with HBSS and incubated with 200 µL of HBSS containing 0.5 µM of compound for 30 to 120 min at 100 rpm. At each time point, medium was sampled before washing the cells with HBSS and extracting intracellular compound using acetonitrile/water (60/40) for 15 min at 500 rpm. Protein content was quantified in representative wells using a ThermoFisher Pierce BCA assay kit. Cellular volume (V_cell_) was calculated assuming 6.5 µL/mg protein (Treyer et al., 2018; Treyer et al., 2019). Experiments were carried out in triplicate on three independent occasions. Compounds were quantified via UPLC-MS. Kp, the intracellular compound accumulation, was calculated according to eq. 1:

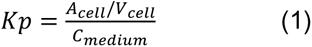

Data were plotted using GraphPad Prism 9.

Binding to lipid-coated beads (f_u,lipid_) was measured with TRANSIL^XL^ Intestinal Absorption Kits (TMP-0100-2096, Sovicell, Leipzig, Germany), as outlined in (Treyer et al., 2019). Briefly, phosphatidylcholine-coated silica beads and 5 µM drug were incubated for 12 min, with orbital shaking. The beads were centrifuged at 750 rpm for 10 min before sampling from the supernatant. Experiments were carried out in triplicate at three independent occasions. Compounds were quantified by LC-MS/MS. The f_u,lipid_ metric, the fraction of unbound compound, was then calculated as outlined in Treyer et al, 2018 and 2019. D_L_, an optimized dilution factor determined by minimizing the sum of the squared prediction errors (Microsoft Excel, Solver add-in), was used to scale f_u,lipid_ to f_u,cell_, the predicted intracellular fraction of unbound compound, according to eq. 2:

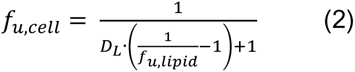

Intracellular bioavailability F_ic_ (Mateus et al., 2017) was calculated from experimentally determined Kp and f_u,cell_ values using eq. 3:

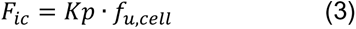

UPLC-MS/MS analysis of sampled fluids was performed on a Waters Acquity UPLC coupled to a Waters Xevo TQ-S micro MS (Milford, MA). Chromatographic separation was achieved using a Waters 1.7 µm C18 BEH column measuring 2 x 50 mm (Milford, MA) with a gradient of 5% to 95% mobile phase B (0.1% formic acid in 100% ACN) in mobile phase A (0.1% formic acid in LC-MS grade water) over a runtime of 2 min. The flow rate was 0.7 mL/min and 7 µL of sample was injected per run. In ESI+ ionization mode, the UPLC-MS parameters listed in Table 2 were used. Data were preprocessed using Waters MassLynx and TargetLynx 4.2.

**Table 2.**
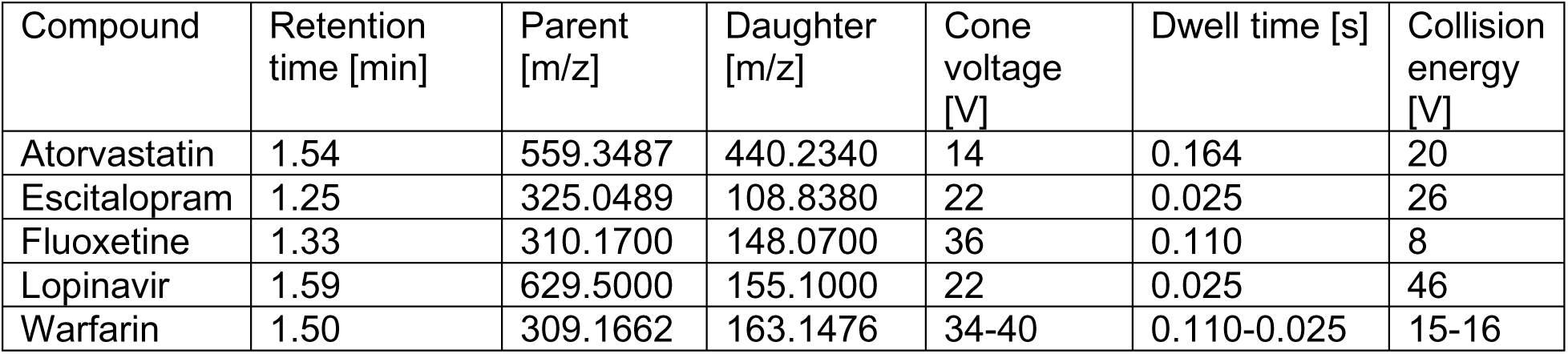
Mass spectrometry parameters for chemical detection of compounds used in HEK cell and lipid-coated bead assays.

| Compound | Retention time [min] | Parent [m/z] | Daughter [m/z] | Cone voltage [V] | Dwell time [s] | Collision energy [V] |
| --- | --- | --- | --- | --- | --- | --- |
| Atorvastatin | 1.54 | 559.3487 | 440.2340 | 14 | 0.164 | 20 |
| Escitalopram | 1.25 | 325.0489 | 108.8380 | 22 | 0.025 | 26 |
| Fluoxetine | 1.33 | 310.1700 | 148.0700 | 36 | 0.110 | 8 |
| Lopinavir | 1.59 | 629.5000 | 155.1000 | 22 | 0.025 | 46 |
| Warfarin | 1.50 | 309.1662 | 163.1476 | 34-40 | 0.110-0.025 | 15-16 |

### Plasmid availability

We will deposit plasmids with the following cDNAs at Addgene:

iFluoxSnFR,

iEscSnFR,

We will deposit the following plasmids at Addgene:

pCMV(MinDis)-iFluoxSnFR_PM,

pCMV(MinDis)-iEscSnFR_PM

pCMV(MinDis)-iFluoxSnFR_cyto

pCMV(MinDis)-iEscSnFR_cyto

pCMV(MinDis)-iFluoxSnFR_ER,

pCMV(MinDis)-iEscSnFR_ER,

pAAV9-hSyn-iFluoxSnFR_PM,

pAAV9-hSyn-iEscSnFR_PM,

pAAV9-hSyn-iFluoxSnFR_cyto,

pAAV9-hSyn iEscSnFR_cyto,

pAAV9-hSyn-iFluoxSnFR_ER,

pAAV9-hSyn-iEscSnFR_ER.

## RESULTS

### Generation of iDrugSnFRs for escitalopram and fluoxetine

To generate iDrugSnFRs for SSRIs, we screened several SSRIs against a panel of biosensors that included our previously published Opu-BC based biosensors (Bera et al., 2019; Borden et al., 2019; Shivange et al., 2019; Unger et al., 2020; Nichols et al., 2022) as well as intermediate constructs from their development process. This screen is described in a protocol under review at Bio-Protocol. From this screen, we identified possible biosensors for fluoxetine and escitalopram. We chose sensors with the lowest EC_50_ for each drug as our starting protein for iDrugSnFR evolution.

We incrementally applied SSM to first- and second-shell amino acid positions within the binding pocket. We evaluated each biosensor and drug partner in lysate from *E. coli* and carried forward the biosensor with the highest S-slope to the subsequent round. S-slope, 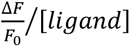 at the beginning of the concentration-response relation, emphasizes the response to ligand concentrations in the pharmacologically relevant range (Bera et al., 2019). Figure 1 summarizes concentration-response relations for the optimized sensors. The escitalopram sensor, iEscSnFR, displayed EC_50_ 4.5 ± 0.2 µM, ΔF_max_/F_0_ 16 ± 0.3, and S-slope 3.6. The fluoxetine sensor, iFluoxSnFR, displayed EC_50_ 8.7 ± 0.2 µM, ΔF_max_/F_0_ 9.2 ± 0.1, and S-slope 1.1. (Fig. 1A).

**Figure 1.**
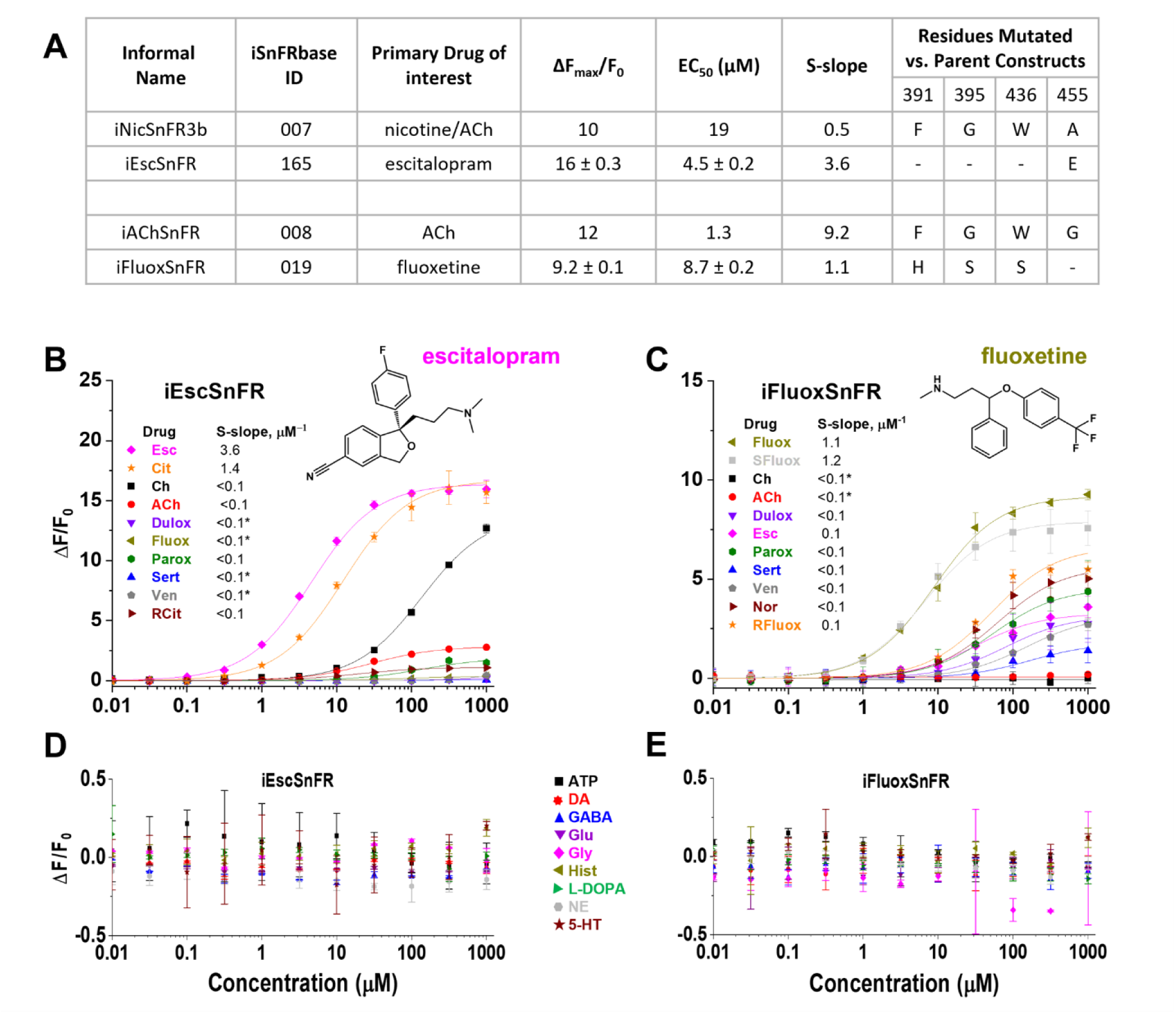
SSRI iDrugSnFR naming, residues mutated, and concentration-response relations. **(A)** Endpoints of SSRI iDrugSnFR development and concentration-response relations versus parent constructs. Data for iAChSnFR from (Borden et al., 2019); Data for iNicSnFR3b from (Shivange et al., 2019) **(B-C)** Concentration-response relations of purified iEscSnFR and iFluoxSnFR versus a drug panel. Abbreviations: Ch choline; ACh, acetylcholine; Dulox, duloxetine; Esc, escitalopram; Fluox, racemic fluoxetine; Parox, paroxetine; Sert, sertraline; Ven, venlafaxine; RCit, R-(-)-citalopram; Cit, racemic citalopram; Nor, norfluoxetine; RFluox, R-(+)- fluoxetine; and SFluox, S-(-)-fluoxetine. Relevant S-slope values for each iDrugSnFR are included in the inset. Dashed lines indicate concentration-response relations that did not approach saturation for the concentration ranges tested; therefore, EC_50_ and ΔF_max_/F0 could not be determined. iEscSnFR **(B)** shows preference for escitalopram over other SSRIs, with measurable binding to choline. **(C)** iFluoxSnFR shows a preference for racemic fluoxetine but also shows modest responses to other SSRIs. (**D**) iEscSnFR and **(E)** iFluoxSnFR shows little or no fluorescence response to all endogenous molecules tested. ATP, adenosine triphosphate; DA, dopamine; GABA, γ-aminobutyric acid; Glu, glutamate; Gly, glycine; Hist, histamine; L-DOPA, levodopa; NE, norepinephrine; 5-HT, serotonin.

### Specificity and thermodynamics of SSRI iDrugSnFRs

We characterized the specificity of purified SSRI iDrugSnFRs for their drug partners versus a panel of related antidepressants, antidepressant metabolites, and nicotinic agonists (Fig. 1B and C). The newly developed iDrugSnFRs showed some sensitivity for other antidepressants. iEscSnFR had greater fidelity for its drug partner, binding few drugs in our panel except for choline (EC_50_ of 140 ± 20 µM, a value ∼10-fold above endogenous levels (Zeisel et al., 1980; Schapiro et al., 1990; Vargas and Jenden, 1996)) (Fig. 1B). iFluoxSnFR showed greater sensitivity for some compounds comprising our drug panel. iFluoxSnFR detected several compounds with EC_50_ values between 32–170 µM, concentrations higher than those relevant for clinical purposes. iFluoxSnFR detected norfluoxetine, the breakdown product of fluoxetine, with an EC_50_ of 63 ± 20 µM, a 9-fold preference in binding for fluoxetine over norfluoxetine. iFluoxSnFR shows no binding to acetylcholine and choline (Fig. 1C). The relative selectivity of each biosensor for its partner compound indicates a structure/function relationship that differs from that of the interaction between hSERT and SSRIs.

We also performed concentration-response experiments with iEscSnFR and iFluoxSnFR against a panel of nine endogenous molecules and their precursors (Fig. 1D and E). Both iEscSnFR and iFluoxSnFR showed no response to any of the nine selected compounds above background.

To examine the thermodynamics of the iDrugSnFR:drug interaction, we conducted ITC binding experiments (Fig. 2A and B). The experimentally determined K_d_ of iEscSnFR, 3.4 ± 0.1 μM, was within a 1.5 factor of the experimentally determined EC_50_ in purified protein (Fig. 2A and B).

**Figure 2.**
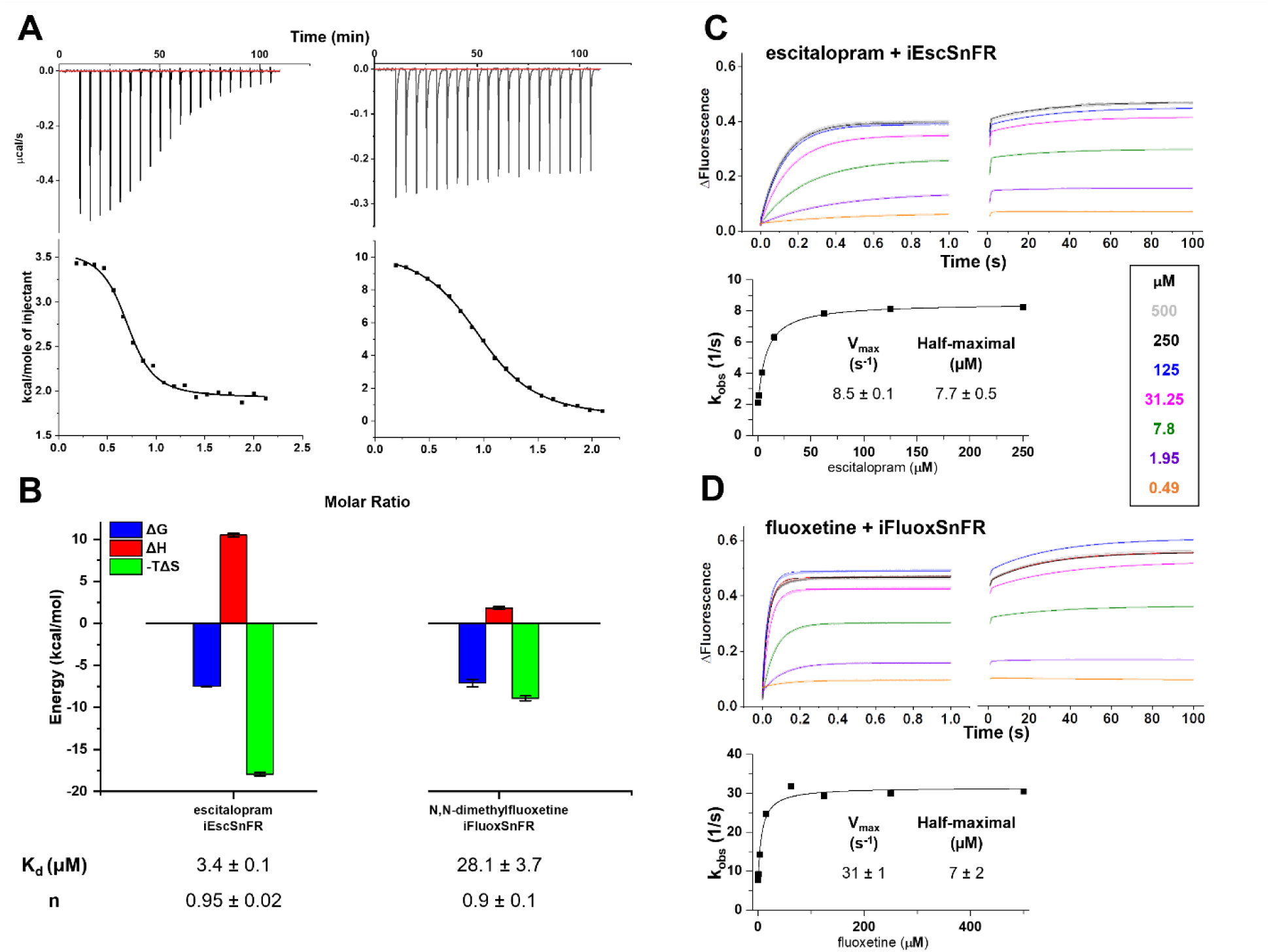
Thermodynamic and kinetic profiles of purified SSRI iDrugSnFR proteins. **(A)** ITC traces and fits. **Top row**: Exemplar heat traces of iEscSnFR paired with escitalopram and iFluoxSnFR paired with N,N-dimethylfluoxetine as obtained by ITC. The heats for iEscSnFR and iFluoxSnFR were endothermic. **Bottom row**: The resulting fits for each iDrugSnFR:drug pair from the integrated heats comprising each series of injections. **(B)** Energy calculations from ITC traces and fits. Both iDrugSnFRs show exergonic reactions, but the relative enthalpic and entropic contributions differ. Affinity (K_D_) and occupancy number (n) were also calculated. Data are from 3 separate runs, Mean ± SEM. Stopped-flow fluorescence data for various concentrations of **(C)** iEscSnFR and **(D)** iFluoxSnFR recorded for periods of 1 and 100 s, at sampling rates of 1 ms and 1 s, respectively. Fluorescence was activated at time zero by mixing agonist and sensor protein as noted. iEscSnFR and iFluoxSnFR data are fits to single exponentials. Plots of the exponential rate constants versus [agonist]s are included for the 1 s data.

When we attempted ITC with iFluoxSnFR and fluoxetine, the low aqueous solubility of fluoxetine led to distortion due to turbulent injections even after multiple attempts with various solvation schemes. Consequently, we performed ITC with N-N-dimethylfluoxetine (Fig. 2A and B). The experiments produced an experimentally determined K_d_ of 28.1 ± 3.7 μM, approximately twice the experimentally determined EC_50_ of iFluoxSnFR for N-N-dimethylfluoxetine (see Fig. 7 below). The ITC data imply that the EC_50_ for fluorescence in iEscSnFR and iFluoxSnFR is dominated by the overall binding of the corresponding ligand.

### Stopped-flow experiments on SSRI iDrugSnFRs

We used a stopped-flow apparatus with millisecond resolution to measure the time course of fluorescent SSRI iDrugSnFRs responses to step-like drug applications (Fig. 2C and D). These data show the trajectory of the ligand-sensor reaction as it relaxes to a new equilibrium after a sudden change in ligand concentration. For both sensors, most of the fluorescence change occurred within the first second with a mono-exponential time course (Fig. 2C and D, upper left panel). An additional, smaller and slower exponential fluorescence increase continued over the next minute (Fig. 2C and D, upper right panel).

In the 1 s stopped-flow experiments, the rate constants for the fluorescence relaxation (k_obs_) were a hyperbolic function of ligand concentration (Fig. 2C and D, lower panels). For escitalopram binding to iEscSnFR, the zero-concentration intercept was 1.8 ± 0.1 s^-1^. The increased k_obs_ was half-maximal at 7.7 ± 0.5 μM escitalopram. We fitted the data to a three-state kinetic mechanism: the apo state, a drug-bound nonfluorescent state, and a rate-limiting conformational change to the fluorescent state. These assumptions predicted an overall steady-state EC_50_ of 1.7 ± 0.3 μM, compared to the value of 4.5 μM obtained with equilibrium concentration-response experiments on iEscSnFR (Fig. 1A). For fluoxetine binding to iFluoxSnFR, the zero-concentration intercept was 6.2 ± 1.5 s^-1^. The increased k_obs_ was half- maximal at 7 ± 2 µM fluoxetine. The 3-state mechanism predicted an overall steady-state EC_50_ of 1.3 ± 0.7 μM, compared to the value of 8.7 μM obtained with equilibrium concentration-response experiments on iFluoxSnFR (Fig. 1A).

We have less complete measurements for the slower phase of the fluorescence increases. The half-maximal amplitudes and rate constants for the slower phases occurred at ligand concentrations in the same concentration range as those for the faster phase. This observation is consistent with the suggestion that the intense excitation beam in the stopped-flow experiments produced further photoactivation of the fluorescent state.

### Characterization of SSRI iDrugSnFRs in primary mouse hippocampal culture

We examined the subcellular pharmacokinetics of the SSRIs in primary mouse hippocampal neurons transduced with AAV vectors encoding the appropriately targeted iDrugSnFRs. The SSRI iDrugSnFRs were targeted to the PM (iDrugSnFR_PM), the endoplasmic reticulum (iDrugSnFR_ER), or the cytoplasm (iDrugSnFR_cyto) as previously described (Bera et al., 2019; Shivange et al., 2019; Nichols et al., 2022). Spinning-disk confocal microscopy showed targeting to the intended organelle or compartment (Fig. 3). ER-targeted biosensor was retained in the ER (Bera et al., 2019; Shivange et al., 2019; Nichols et al., 2022). iDrugSnFR targeted to the PM showed correct localization, with some iDrugSnFR observed in the cell interior (most likely as part of the cellular membrane trafficking system or inclusion bodies). The cytoplasm-targeted constructs appeared in both soma and dendrites.

**Figure 3.**
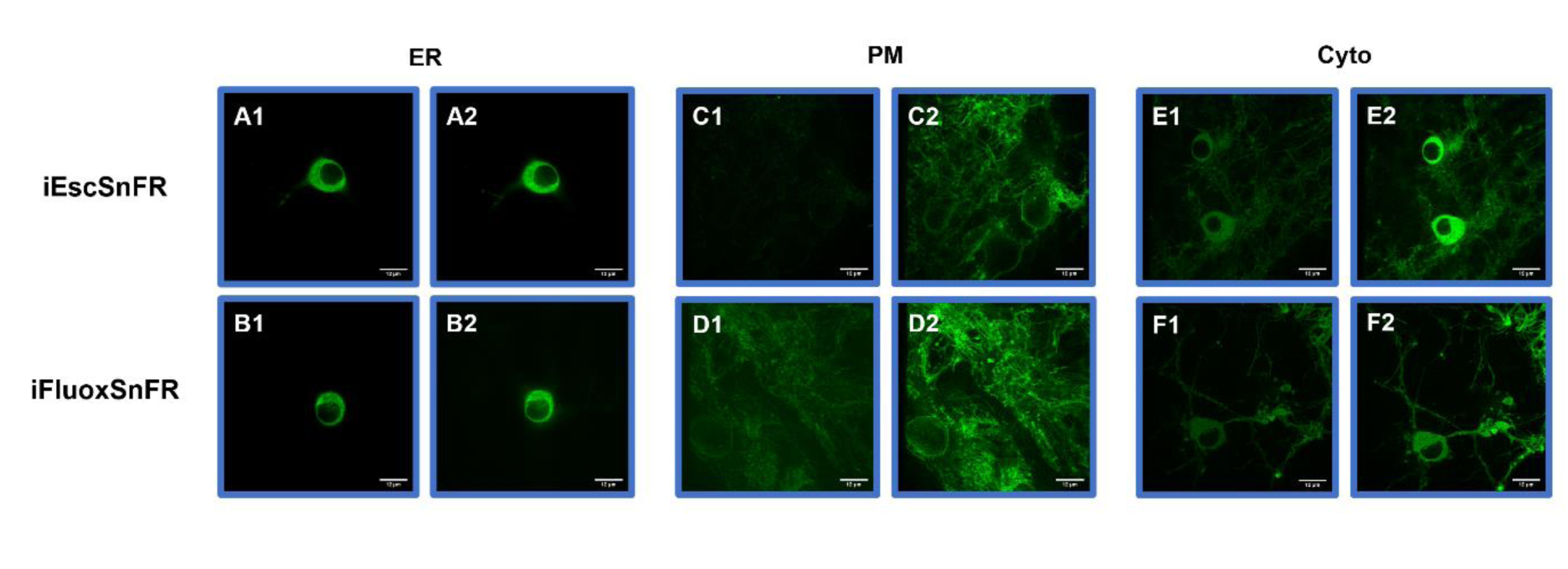
Spinning disk laser scanning confocal inverted microscope images of SSRI iDrugSnFRs in primary mouse hippocampal neurons. ER-targeted constructs of iEscSnFR and iFluoxSnFR are shown before **(A1-B1)** and during **(A2-B2)** exposure to each drug partner at 10 μM. ER-targeted iDrugSnFRs show the eponymous reticulated pattern, and fluorescence is excluded from the nucleus. PM-targeted constructs of the same iDrugSnFRs are shown before **(C1-D1)** and after **(C2-D2)** drug introduction. Localization in the PM is robust, with some minimal puncta that may represent inclusion bodies or internal transport. Cyto-targeted constructs of iEscSnFR and iFluoxSnFR are shown before **(E1-F1)** and after **(E2-F2)** exposure to each drug partner.

We then performed concentration-response experiments in primary mouse hippocampal culture using wide-field fluorescence imaging with each iDrugSnFR and its drug partner, sampling a range of concentrations approximately an order of magnitude above and below the EC_50_ as determined for the purified protein (Fig. 4, Multimedia files 1-4). iEscSnFR showed a robust response to escitalopram at the PM and the ER across a range of concentrations from 0.1‒31.6 µM, and the speed was nearly limited by solution exchanges; there was a clear return to baseline fluorescence after each drug application on the order of seconds (Fig. 4A). A maximum ΔF/F_0_ value of ∼ 2 was reached at 31.6 μM with iEscSnFR_ER construct. We also observed a 10% higher ΔF/F_0_ in the _ER construct versus the _PM construct in concentrations above 1 μM, a phenomenon we had not encountered in any of our previous work.

**Figure 4.**
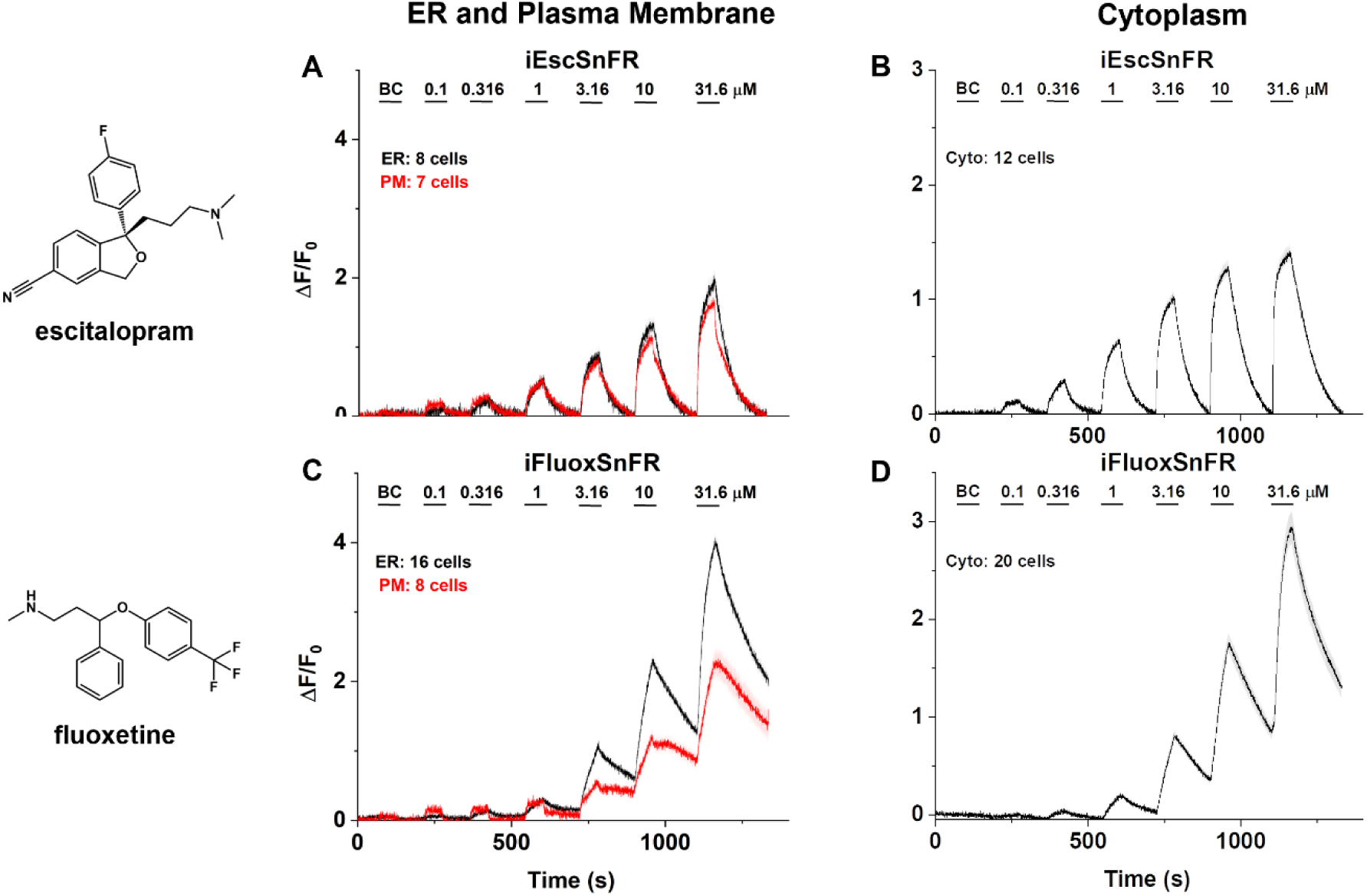
SSRI iDrugSnFR concentration-response relations in primary hippocampal culture. **(A-D)** Each iDrugSnFR detects its drug partner at the endoplasmic reticulum (ER), plasma membrane (PM), or cytoplasm (cyto) of primary hippocampal culture at the concentrations sampled. BC = Buffer control. SEM of data are indicated by semi-transparent shrouds around traces where trace width is exceeded. Drugs were applied for 60 s pulses at 120–150 s intervals. **(A-B)** iEscSnFR detects escitalopram, approaching a plateau during the application, then, returns to baseline fluorescence during the washout, at all targeted locations. **(C-D)** iFluoxSnFR detection of fluoxetine has not yet reached a plateau during the application, then shows an incomplete washout with no return to baseline fluorescence during the washout period, in every targeted location.

In contrast, both iFluoxSnFR_ER and iFluoxSnFR_PM constructs detected fluoxetine across a range of concentrations, but after 60 s of drug application the ΔF/F_0_ had not begun to plateau to a maximum value at concentrations of 1 μM and higher (Fig. 4C). Therefore, responses to fluoxetine increases and decreases were much slower than solution changes for both constructs, on the order of hundreds of s. We also observed that the ΔF/F_0_ of iFluoxSnFR_ER was ∼2-fold higher than iFluoxSnFR_PM at concentrations 3.16 μM and higher.

Concentration-response experiments in primary hippocampal culture with iDrugSnFR_cyto constructs demonstrated washout dynamics for escitalopram and fluoxetine similar to those obtained when the sensor was targeted to the PM and ER (Fig. 4B and D).

To further examine the extended kinetics of fluoxetine we observed with iFluoxSnFR, we recorded the fluorescence waveforms for fluoxetine at 1 μM with an extended application time of 10 min and a washout time of 12 min (Fig. 5A). For the PM, the kinetics clearly showed two components. The faster component represented ∼10% of the total change and is indistinguishable from the solution change. The slower component had time constants of 200-300 s for both the wash-in and washout in both iFluoxSnFR_PM and iFluoxSnFR_ER. After a 12 min washout, both the iFluoxSnFR_PM and _ER construct neared baseline fluorescence, indicating that full washout of fluoxetine can be achieved, but on a time scale nearly a log unit slower than for drugs such as nicotine and ketamine, and two times slower than cytisine (Bera et al., 2019; Shivange et al., 2019; Muthusamy et al., 2022; Nichols et al., 2022).

**Figure 5.**
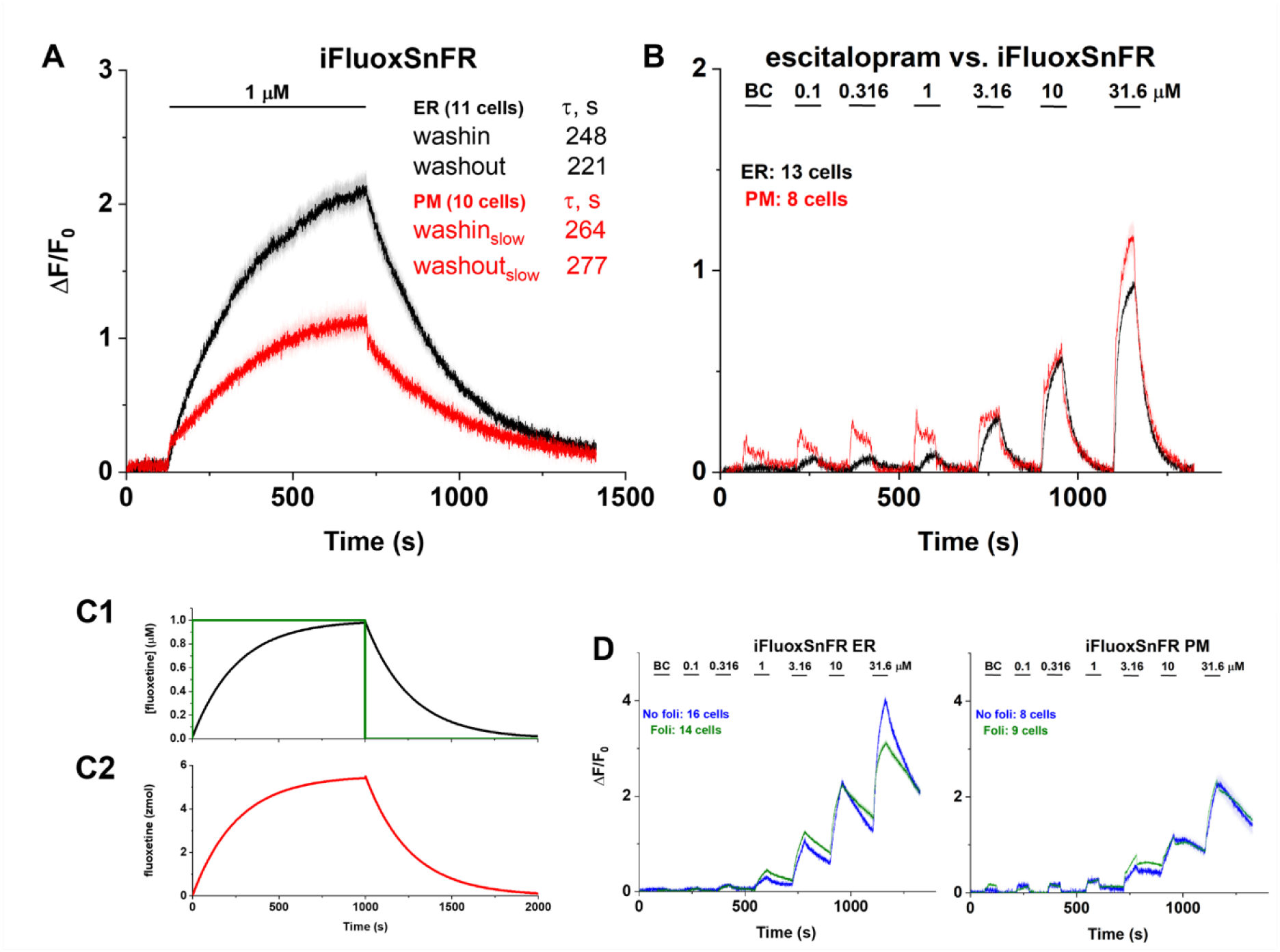
Further analysis of fluoxetine kinetics in primary hippocampal neurons. **(A)** Traces of fluorescence responses during exposure to 1 μM fluoxetine with iFluoxSnFR. BC = Buffer control. SEM of data are indicated by semi-transparent shrouds around traces where trace width is exceeded. Marks show t_1/2_ (half-rise time). A relatively long application (600 s) allowed ER- and PM-targeted iFluoxSnFR detection of 1 μM fluoxetine to approach a maximum ΔF/F_0_. A slightly longer (720 s) washout allowed a return to baseline fluorescence for both ER- and PM- targeted iFluoxSnFR. **(B)** A control experiment: imaging concentration-response relations for escitalopram against iFluoxSnFR. BC = Buffer control. SEM of data are indicated by semi- transparent shrouds around traces where trace width is exceeded. iFluoxSnFR detects escitalopram at both the PM and ER. Escitalopram enters and exits the ER with a return to baseline fluorescence during the washout, a direct contrast to the behavior of fluoxetine as detected by iFluoxSnFR. **(C)** Simulations of fluoxetine in the extracellular space, plasma membrane, and cytoplasm of a spherical cell. C1, the green trace gives the applied (“clamped”) [fluoxetine] in a shell 11.5 μm from the center of the cell. At a radius of 11.5 μm in the extracellular solution, the concentration is stepped from zero to 1 μM for 1000 s; the concentration is then stepped back to zero (green trace). The concentrations in all extracellular shells (between the 11.5 μm shell and the PM shell at 7.5 μm radius) equilibrate within ∼ 50 ms and are indistinguishable from the applied concentration on this time scale. The black trace gives the cytoplasmic [fluoxetine] within the shell of outer radius of 7.495 μm, 10 nm below the plasma membrane. The concentrations in all other intracellular shells show a dispersion of ∼ 50 ms and are indistinguishable from the black trace on this time scale. The intracellular [fluoxetine] resembles that of panel A. C2, the moles of fluoxetine bound within the simulated “membrane shell”. With the parameters given in Table 1, the time course of PM-bound fluoxetine is indistinguishable from that of intracellular [fluoxetine] and resembles that of panel A. See Methods, text, and Table 1. **(D)** Pretreatment of primary hippocampal neurons with 80 nM folimycin does not substantially alter the concentration-response relations for iFluoxSnFR against fluoxetine versus untreated neurons in a side-by-side experiment.

To confirm that the iFluoxSnFR_PM and _ER constructs functioned as expected, and to ensure that the extended kinetics of fluoxetine we observed in primary hippocampal culture was not the result of idiosyncratic biosensor function or folding in neurons, we tested iFluoxSnFR_PM and iFluoxSnFR_ER versus escitalopram. iFluoxSnFR binds escitalopram in the same concentration range as fluoxetine (though right shifted and with lower ΔF/F_0_) (Fig. 1C). After viral transduction of iFluoxSnFR_PM and iFluoxSnFR_ER, we performed time-resolved imaging for pulses of 0.1–31.6 μM escitalopram (Fig. 5B). These escitalopram waveforms resembled those of iEscSnFR detection of escitalopram in primary mouse hippocampal culture (Fig. 4A), confirming that iFluoxSnFR_ER and _PM function as expected. Thus, the slower kinetics for iFluoxSnFR_ER and iFluoxSnFR_PM arise from a property inherent to the interaction between fluoxetine and the primary hippocampal culture.

### Estimating fluoxetine accumulation in the neuronal membrane

That the fluoxetine signals show time constants of 200-300 s at all three locations (plasma membrane, ER, and cytoplasm) led us to suspect the existence of a local binding site(s) that delays the appearance and disappearance of fluoxetine near neurons. A related phenomenon is termed “buffered diffusion” (Armstrong and Lester, 1979).

We based our analysis on the unusually high pharmacokinetically defined volume of distribution exhibited by all SSRIs (see Introduction). Basic compounds can accumulate within the body via two major mechanisms: acid trapping within low-pH organelles and drug partitioning into membrane lipids (Smith et al., 2012). Acid trapping within low-pH organelles has been suggested for nicotine, antipsychotics, and ketamine (Lester et al., 2009; Tischbirek et al., 2012; Lester et al., 2015; Tucker et al., 2015; Govind et al., 2017). Therefore, we first tested for vesicular accumulation by blocking the vesicular proton pump with folimycin. We found little to no effect of such blockade (Fig. 5D).

We therefore turned our attention to drug partitioning into membrane lipids. Membrane partitioning should be distinguished from the more familiar, readily modelled fact that the protonated, charged species permeates membranes much more slowly than the uncharged, neutral species. Because both escitalopram and fluoxetine have calculated pKa ∼ 9.8, the charge-dominated effect is expected to decrease the effective diffusion constant (n_pH_) by at least two orders of magnitude (Yu et al., 2016).

Recent studies show how membrane partitioning of basic molecules plays a role in some molecular and cellular bases of the classical volume of distribution (Loryan et al., 2013; Mateus et al., 2014; Treyer et al., 2019). The nitrogen interacts with the phospholipid head groups while the less polar moieties interact with the fatty acid tails (Mateus et al., 2014; Kapoor et al., 2019). The equilibrium parameters of such accumulation have been estimated by direct measurements on membrane-coated beads (see also below), by ITC, and by perturbation of gramicidin gating (Kapoor et al., 2019; Treyer et al., 2019). However, the kinetics of this accumulation are relatively unstudied and may be revealed for the first time by measurements such as Figures 4C, 4D and 5A.

The iFluoxSnFR measurements did show that locally measured SSRI concentrations eventually reach the applied concentration; the novel observation is that the approach to steady state at the PM required several hundred s. We were able to simulate these delays (Fig. 5C) only by assuming that the extracellular facing iFluoxSnFR_PM measures, at least partially, the membrane-bound fluoxetine as it increases or decreases in response to step changes in the externally applied solutions. For previously reported PM-anchored iDrugSnFRs, the measurements were dominated by the free concentration in the extracellular aqueous phase (Bera et al., 2019; Shivange et al., 2019; Muthusamy et al., 2022; Nichols et al., 2022). Similarly, for membrane-excluded quaternary SSRI derivatives, the PM-anchored SSRI iDrugSnFRs also measured the aqueous concentrations of the drugs (Figs. 8 and 9 below). We suggest that the unique signals produced by fluoxetine at PM-localized iFluoxSnFR arise from two facts. First, if the anomalously high volume of distribution arises from the membrane accumulation, then this accumulation exceeds the aqueous concentration by orders of magnitude (the next paragraph gives an estimate). Second, PBPs from bacteria and archaea are specialized to transfer the ligand directly to membrane-embedded transporters that are adjacent (within just a few Å) to their PBP binding site (Scheepers et al., 2016; Nguyen et al., 2018) (also PDB entries 2ONK, 4TQU, 2R6G, 6CVL, 4FI3, 5B58), in contrast to the several μm thick unstirred layer inferred from ITC measurements on the fluoxetine-lipid interaction (Kapoor et al., 2019).

Our data yielded an estimate of membrane partitioning. With the unique sensing assumption discussed above, we modeled the fluoxetine measurements by assuming that the effective diffusion coefficient is reduced further by lipid binding within the membrane (Crank, 1975). Table 1 gives our assumptions for the better-characterized underlying parameters. The most important adjustable parameter is the binding constant K_d_ for lipid-fluoxetine binding. The only available measurement is “at least 100 μM” (Mateus et al., 2013). Because we treat membrane permeation as a single first-order process whose kinetics are orders of magnitude slower than diffusion in the cytoplasm and extracellular solution, the simulation predicts exponential kinetics. The experimentally measured time constant of 200-300 s (Fig. 5A) was explained by a K_d_ of 2.2 mM. The extent of membrane accumulation is therefore (lipid molarity in the shell)/(fluoxetine-lipid K_d_), or 181-fold higher than the free solution value of fluoxetine.

### Characterization of SSRI iDrugSnFRs in HeLa cells

In light of the surprisingly slow kinetics from imaging of iFluoxSnFR movements in primary cultured neurons, we examined the subcellular pharmacokinetics of the SSRIs in a transfected mammalian cell line. The SSRI iDrugSnFRs were targeted to the PM (iDrugSnFR_PM), or the ER (iDrugSnFR_ER) as previously described (Bera et al., 2019; Shivange et al., 2019; Muthusamy et al., 2022; Nichols et al., 2022). We also assembled iEscSnFR and iFluoxSnFR constructs targeted to the cytoplasm (iEscSnFR_cyto and iFluoxSnFR_cyto) for use in HeLa cells. To examine the localization of the three constructs at higher optical resolution, we imaged HeLa cell cultures using a spinning disk laser scanning inverted confocal microscope (Fig. 6A-C, F-H). Localization of the biosensor resembled previously described iDrugSnFR _PM and _ER constructs (Bera et al., 2019; Shivange et al., 2019; Muthusamy et al., 2022; Nichols et al., 2022) and the localization pattern of iEscSnFR and iFluoxSnFR in primary hippocampal culture imaging (Fig. 3). The _cyto construct was excluded from the nucleus but otherwise showed a relatively featureless intracellular pattern.

**Figure 6.**
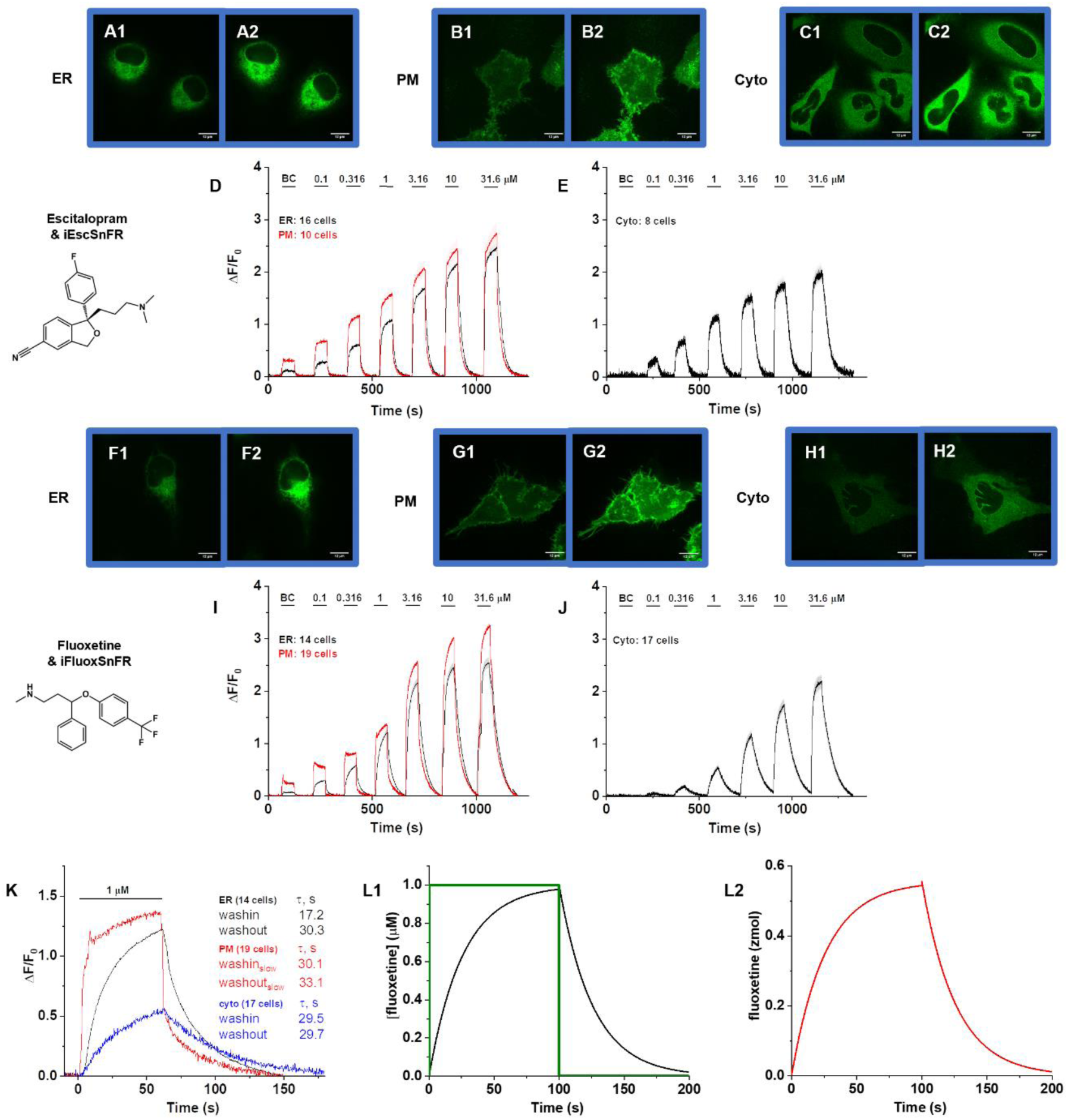
Spinning disk laser scanning confocal inverted microscope images of SSRI iDrugSnFRs and ER-, PM-, and cytoplasm-targeted SSRI iDrugSnFR concentration- response relations in HeLa cells. **(A-C, F-G)** ER-targeted constructs of iEscSnFR and iFluoxSnFR are shown before **(A1, F1)** and during **(A2, F2)** exposure. ER-targeted iDrugSnFRs show the eponymous reticulated structure and dark ovals corresponding to the nucleus. PM- targeted constructs of both SSRI iDrugSnFRs are shown before **(B1, G1)** and after **(B2, G2)** drug introduction. Localization to the PM is robust, with some minimal puncta that may represent inclusion bodies or internal transport. Cytoplasm-targeted constructs of iEscSnFR and iFluoxSnFR are shown before **(C1, H1)** and after **(C2, H2)** exposure to each drug partner at 10 μM. **(D-E, I-J)** Drugs were applied for 60 s pulses at 90–120 s intervals. Each iDrugSnFR detects its drug partner at the PM, ER, and cytoplasm of HeLa cells at the concentrations sampled. BC = Buffer control. SEM of data are indicated by semi-transparent shrouds around traces where trace width is exceeded. **(D-E)** iEscSnFR detects escitalopram, approaching a plateau during the application, then returns to baseline fluorescence during the washout, when targeted to the ER, PM, and cytoplasm. **(I)** iFluoxSnFR targeted to the ER and PM detects fluoxetine with a return to baseline fluorescence between applications. **(J)** iFluoxSnFR targeted to the cytoplasm detects fluoxetine with a return to baseline fluorescence between applications. **(K)** Superimposed waveforms for a 60 s pulse of 1 μM fluoxetine vs. iFluoxSnFR targeted to the ER, PM, and cytoplasm in HeLa cells. Tabular values give the time constants of each phase for ER and cytoplasm as well as the time constants for the slower phase for the PM. **(L)** Simulations of the [fluoxetine] within intracellular shells. All intracellular shells superimpose on this time scale. The green and black traces are equivalent to their counterpart in Fig. 5C except that we have presumed weaker membrane accumulation than in the hippocampal neuron PM (L1). L2, simulated accumulation of fluoxetine within the simulated “membrane shell”, corresponding to the slower phase of panel K for the PM-localized sensor.

We performed imaging concentration-response experiments in HeLa cells using wide-field fluorescence imaging with each iDrugSnFR and its drug partner, applying the same concentrations as in the neuronal cell culture experiments (Fig. 6D-E and I-J). Compared with the cultured neuron experiments, the HeLa cell experiments showed larger ΔF/F_0_ across all concentrations sampled for the _PM, _ER, and _cyto constructs (Fig. 4), primarily because the very thin HeLa cells have little endogenous fluorescence and therefore comparatively small F_0_.

iEscSnFR showed a robust response to escitalopram at the PM, ER, and cytoplasm of HeLa cells across a range of concentrations from 0.1‒31.6 µM, and the speed was nearly limited by solution exchanges. At 31.6 µM, the PM had ΔF/F_0_ of ∼2.75, while the ER had ΔF/F_0_ of ∼2.5; at concentrations below this value, the ER had ∼30–80% of the PM signal, which indicated a difference in membrane crossing (Fig. 6D). The _cyto construct had a maximal ΔF/F_0_ of ∼2 at 31.6 µM.

The iFluoxSnFR_PM construct detected fluoxetine across a range of concentrations, reaching a maximum ΔF/F_0_ of ∼3.25 at 31.6 µM, with the _ER construct displaying ∼50–80% of the signal seen in the PM construct (Fig. 6I). The _cyto construct had a maximal ΔF/F_0_ of ∼2.25 at 31.6 µM. iFluoxSnFR targeted to the PM, cytoplasm, or ER in HeLa cells showed wash-in and washout kinetics characteristics that were slower than the solution changes but ∼10-fold more rapid than in hippocampal cultures. At 1 μM fluoxetine (Fig. 6K), the _ER and _cyto constructs displayed single exponential kinetics, as in the neuronal cultures. The iFluoxSnFR_PM construct showed two phases during the wash-in and washout, like the same construct expressed in neurons. As in neurons, the faster phase was indistinguishable from the solution change; but it accounted for ∼80% of the waveform, in contrast to the ∼10% in neurons.

We simulated the slower phase of fluoxetine kinetics in HeLa cells using the diffusion-binding model (Fig. 6L). We assumed that fluoxetine accumulation in the membrane is governed by a fluoxetine-lipid K_d_ of 22 mM, or ∼ 10-fold weaker than in hippocampal neurons. This assumption of weaker membrane accumulation may also explain how iFluoxSnFR_PM signal is dominated by the [fluoxetine] in the extracellular solution, with only a small contribution from fluoxetine accumulated in the PM.

### Cellular Experiments with impermeant SSRI derivatives

We performed concentration-response relations for purified iEscSnFR and iFluoxSnFR with the quaternary derivatives (Fig. 7). The ΔF/F_0_ of iEscSnFR with N-methylescitalopram and escitalopram was nearly identical at ∼16, but iEscSnFR had an approximately 2-fold lower EC_50_ for N-methylescitalopram at 1.8 ± 0.2 µM (Fig. 7A). iFluoxSnFR detected N-N-dimethylfluoxetine with ΔF/F_0_ of 5.0 ± 0.1, which was lower than that for fluoxetine (6.6 ± 0.1). The EC_50_ of iFluoxSnFR for N-N-dimethylfluoxetine was 14 ± 0.4 versus the EC_50_ of 8.3 ± 0.6 µM, an approximate two-fold shift in affinity (Fig. 7B).

**Figure 7.**
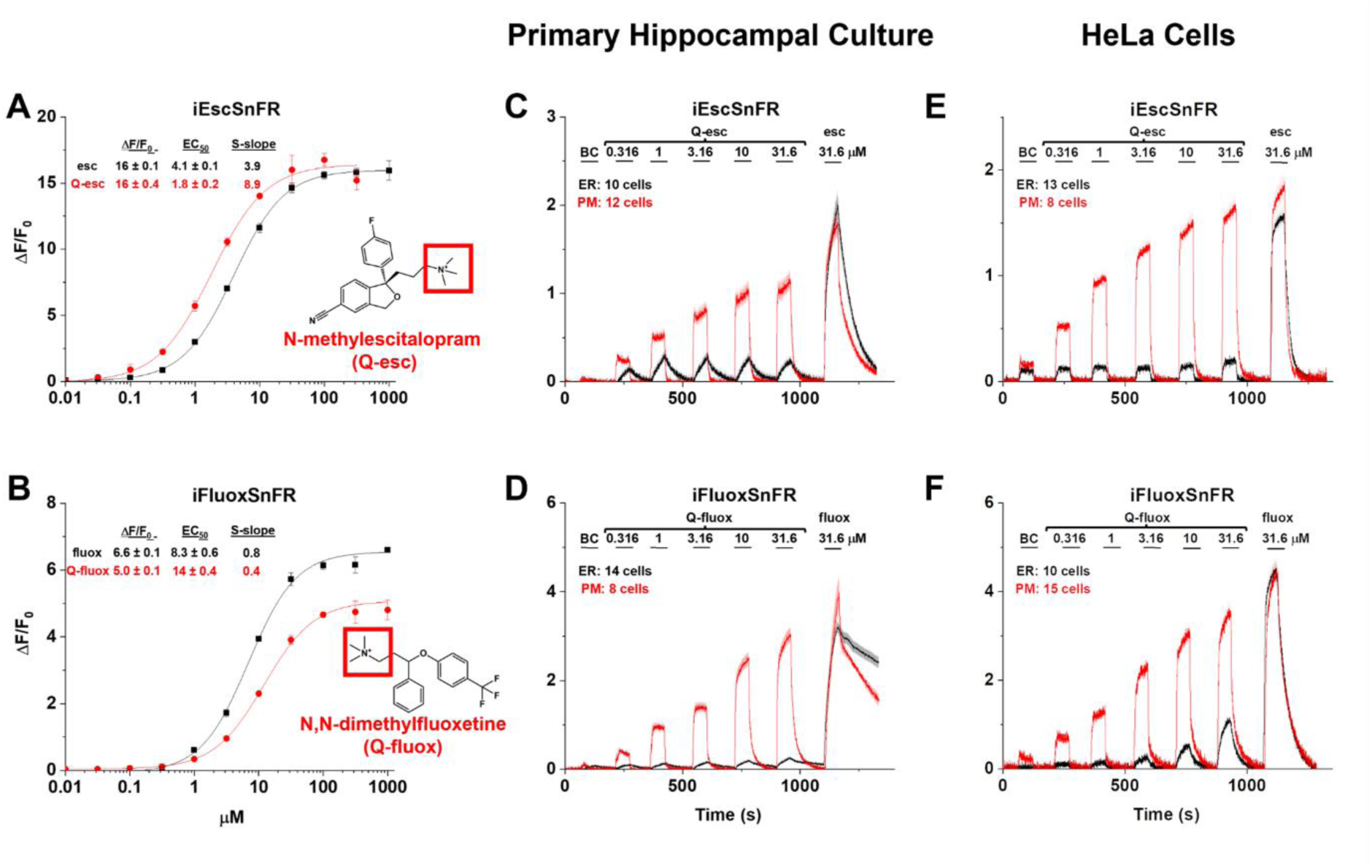
Quaternary SSRI derivatives: SSRI iDrugSnFR concentration-response relations in purified protein, primary hippocampal culture, and HeLa cells. **(A, B)** *In vitro* dose– response relations of purified SSRI iDrugSnFRs against quaternary SSRI derivatives. Abbreviations: esc, escitalopram; Q-esc, N-methylescitalopram; fluox, racemic fluoxetine; Q- fluox, N,N-dimethylfluoxetine. **(A)** iEscSnFR detects N-methylescitalopram with an EC_50_ ∼ half that for escitalopram. **(B)** iFluoxSnFR detects N,N-dimethylfluoxetine with an EC_50_ ∼ twice that for fluoxetine. **(C-F)** Each iDrugSnFR detects its drug partner at the concentrations sampled in primary hippocampal culture and HeLa cells. BC = Buffer control. SEM of data are indicated by semi-transparent shrouds around traces where trace width is exceeded. **(C, E)** Drugs were applied for 60 s pulses at 90–120 s intervals to cells expressing _ER or _PM constructs. In these data, iEscSnFR_PM detects the presence of N-methylescitalopram with a near approach to a plateau during the application, with a return to baseline fluorescence during the washout. In contrast, iEscSnFR_ER is unable to detect N-methylescitalopram. A control dose of escitalopram (final application) is detected by both the PM and ER-targeted constructs. **(D, F)** In cellular experiments, iFluoxSnFR_PM detects fluoxetine with a near approach to a plateau during the application, with a return to baseline during the washout. In contrast, iFluoxSnFR_ER in primary hippocampal culture does not detect N,N-dimethylfluoxetine and iFluoxSnFR_ER in HeLa cells only detects N,N-dimethylfluoxetine above BC only at concentrations above 10 μM. A control dose of fluoxetine is detected by both the PM and ER-targeted constructs (final application). Application of fluoxetine in primary hippocampal culture reproduces the slowly increasing rising phase and the extended washout observed in the experiment of Figure 4.

In concentration-response experiments with quaternary SSRIs in primary mouse hippocampal culture, the speed of the wash-in and washout phases was nearly limited by solution exchanges for both iEscSnFR_PM and iFluoxSnFR_PM (Fig. 7C and D). The application of 31.6 µM SSRI following the quaternary SSRI dosing (designed to act as a control) exhibited a kinetic profile similar to the equivalent concentration in previous concentration-response experiments in primary mouse hippocampal culture (Fig. 4A and C). Of particular note, the kinetic profile of N,N- dimethylfluoxetine as detected by iFluoxSnFR_PM showed a return to baseline fluorescence within seconds after drug washout, a distinctly different result from the observed profile of fluoxetine as detected by iFluoxSnFR_PM. iEscSnFR_ER and iFluoxSnFR_ER showed little ΔF/F_0_ response to application of their corresponding quaternary derivatives, presumably because the permanent positive charges on the quaternary drugs result in a reduced ability to cross membranes.

We also performed concentration-response experiments with the quaternary SSRIs in HeLa cells transfected with PM- and ER-targeted constructs of iEscSnFR and iFluoxSnFR (Fig. 7E and F). The PM-targeted constructs detected their respective quaternary SSRI derivatives over the 0.1–31.6 µM range sampled, with characteristics similar to those detected in primary hippocampal culture (Fig. 7C and D). The detection of quaternary SSRI by the ER-targeted constructs was likewise minimal, with the exception that iFluoxSnFR_ER had ΔF/F_0_ above baseline for N-N-dimethylfluoxetine at concentrations above 3.16 µM. The iFluoxSnFR_ER signal above baseline stays below ∼20% of the fluorescence signal of the _PM construct and represents concentrations above clinical relevance.

To examine the limits of membrane impermeability for quaternary SSRI derivatives, we tested an extended period of co-incubation (Fig. 8). We transfected the _ER and _PM constructs of both iEscSnFR and iFluoxSnFR into HeLa cells and incubated these cells with 500 nM drug (a concentration with appreciable ΔF/F_0_ and within a log unit of the physiologically relevant concentrations of escitalopram and fluoxetine *in vivo* (Karson et al., 1992; Renshaw et al., 1992; Bolo et al., 2000; Paulzen et al., 2016)). We incubated transfected HeLa cells with either an SSRI or a quaternary derivative for 2.4 h (Fig. 8). After transfer to the imaging rig and an equilibration period with buffer containing an identical concentration of the incubation drug, we started a program that included a buffer wash, a short introduction of the complementary compound (i.e. quaternary SSRI if the incubation drug was an SSRI or vice versa), a second buffer wash, and finally a reintroduction of the incubation compound (Fig. 8).

**Figure 8.**
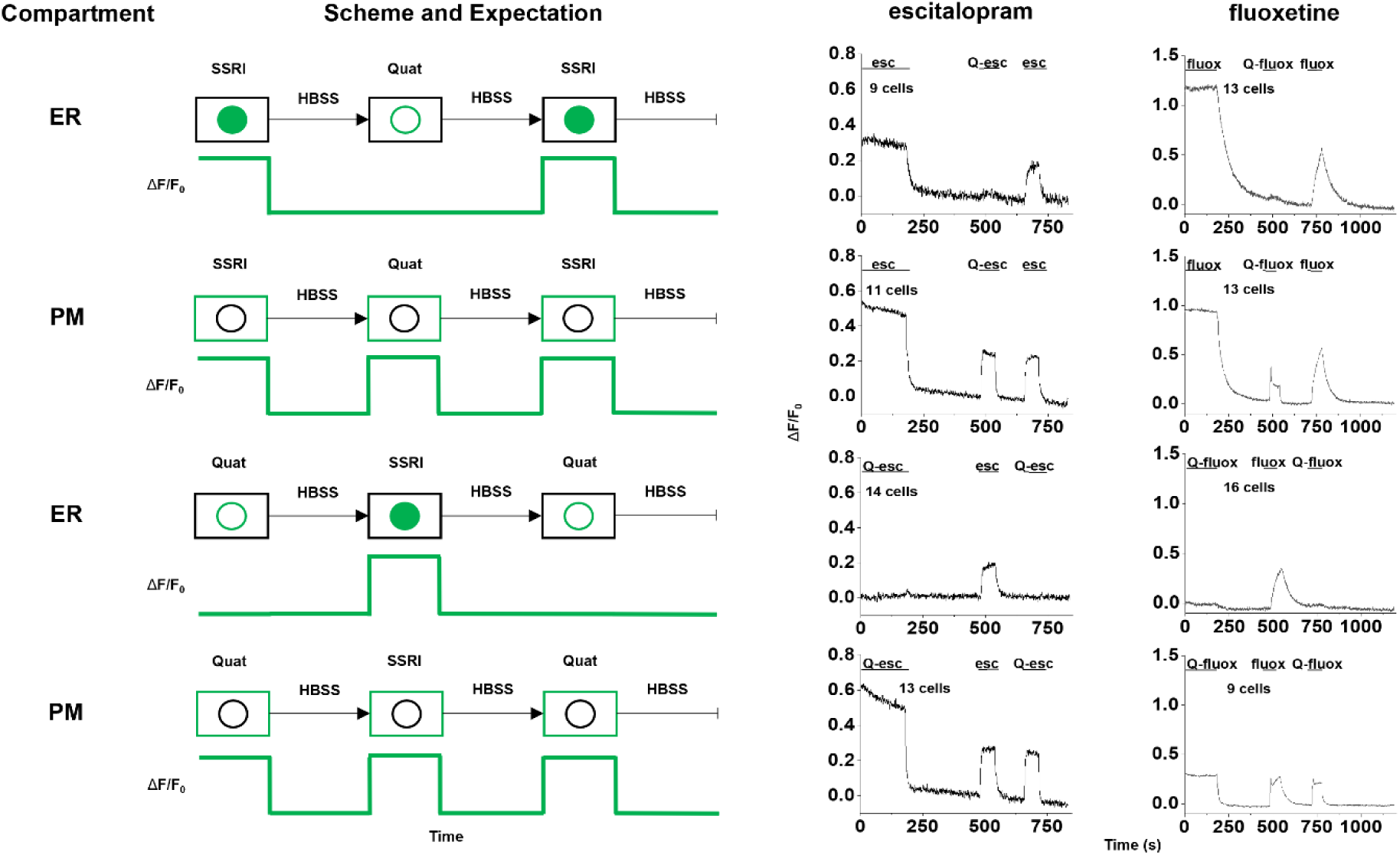
2.4-hour incubation of SSRIs and quaternary derivatives with HeLa cells. Abbreviations: esc, escitalopram; Q-esc, N-methylescitalopram; fluox, racemic fluoxetine; Q- fluox, N,N-dimethylfluoxetine. **Left column**: Targeted compartment of the SSRI biosensor. **Middle column**: Scheme and expectation of fluorescence response by biosensor based on compartment targeted and pre-incubated drug. Following pre-incubation, the drug is washed out, after which the alternate drug is washed in (i.e. when SSRI was pre-incubated, the quaternary derivative was applied and vice versa). An additional washout follows; then the originally pre- incubated drug is reapplied. **Right columns**: Fluorescence response of escitalopram and fluoxetine by their corresponding iDrugSnFR after pre-incubation, washes, and subsequent drug applications, agreeing with the expectations described for the middle column.

**Figure 9.**
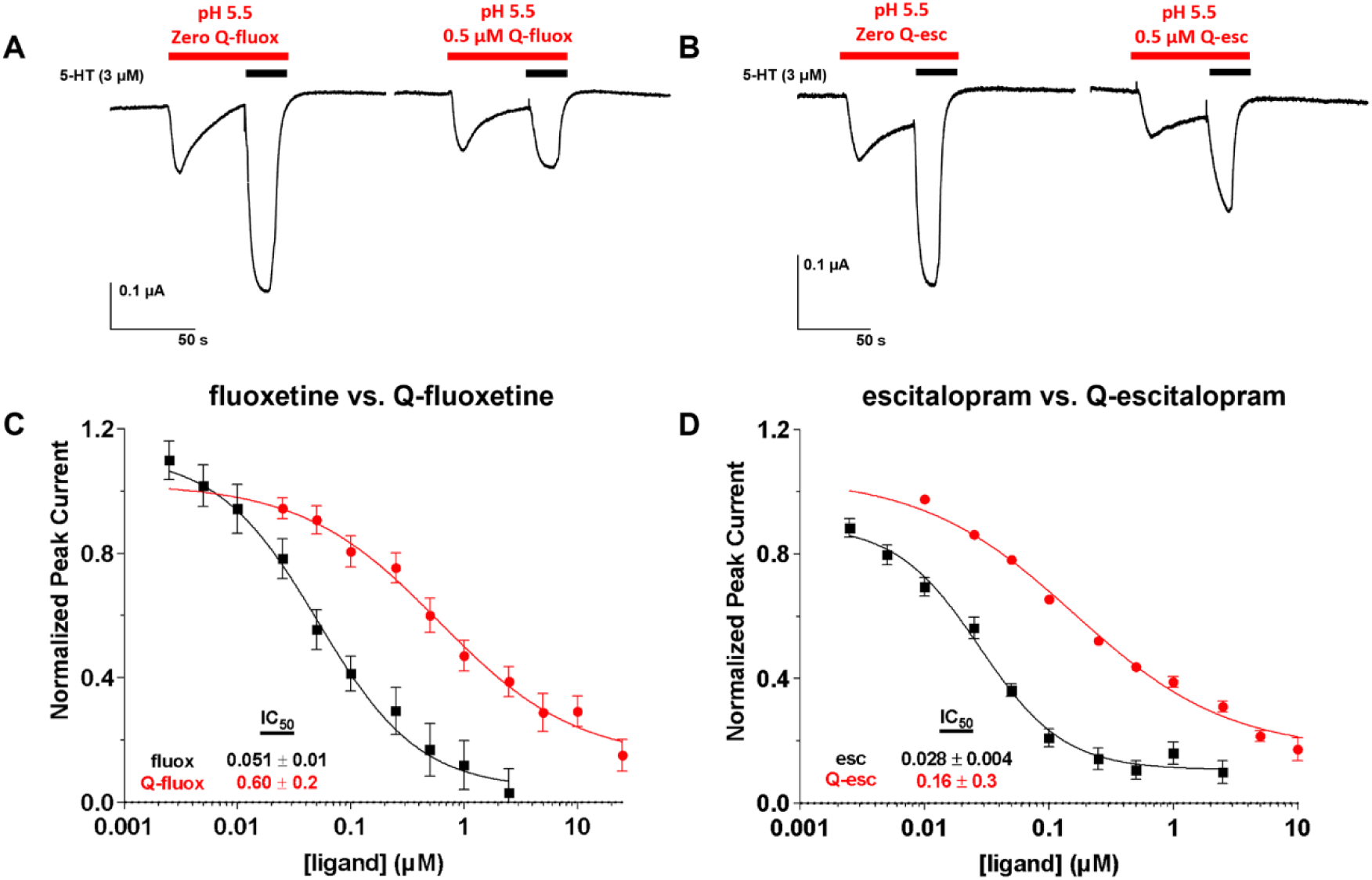
Inhibition of 5-HT induced hSERT transport-associated currents by SSRIs and their quaternary derivatives. Abbreviations: Esc, escitalopram; Q-esc, N-methylescitalopram; fluox, racemic fluoxetine; Q-fluox, N,N-dimethylfluoxetine. **(A-B)** Exemplar traces of 5-HT-induced hSERT currents in the absence and presence of Q-fluox and Q-Esc respectively. **(C-D)** Inhibition of 5-HT-induced hSERT currents of SSRIs and quaternary SSRIs derivatives fluoxetine and escitalopram respectively. IC_50_ values and Hill coefficient calculated from the corresponding fit. **(C)** N,N-dimethylfluoxetine (n = 11) had an IC_50_ 12-fold higher than fluoxetine (n = 13) for the inhibition of hSERT transport-associated currents. **(D)** N-methylescitalopram (n =24) had an IC_50_ 6-fold higher than escitalopram (n =18) for the inhibition of hSERT transport-associated currents.

When SSRIs were pre-incubated with _ER constructs, we saw an initial fluorescence signal that indicated that the SSRIs were present in the ER. Application of buffer decreased the fluorescence signal to a new baseline. Subsequent application of the quaternary SSRIs caused no appreciable increase in fluorescence signal, presumably because the quaternary SSRIs were unable to cross into the ER. A reapplication of the SSRIs also provided biosensor fluorescence signal over background in the ER, though the ΔF/F_0_ of the reapplication is ∼50% of the signal observed after the 2.4 h incubation. Possibly the ΔF/F_0_ would have returned to its maximum value if the reapplication occurred over a longer period (Fig. 8, first row).

When the SSRIs were pre-incubated with cells expressing the _PM construct, introduction of control HBSS (Fig. 8, second row) produced a decrease to a new baseline. As would be predicted, in this case, reapplication of quaternary SSRIs generated a reversible fluorescence increase, because PM-targeted biosensor was accessible to detect the quaternary drug. Reapplication of the SSRIs once again generated a fluorescence signal over baseline, though once again, this signal was ∼50% of the signal inferred from the end of the 2.4 h incubation (Fig. 8, second row).

When quaternary SSRIs were pre-incubated with ER-targeted biosensors, introduction of control HBSS (Fig. 8, third row) did not produce a clear decrease in biosensor fluorescence signal. Rather, the signal we observed continued as the existing baseline, with little to no change in signal. Upon application of SSRI, we observed a clear reversible increase in ΔF/F_0_ over the baseline fluorescence signal in the ER, which indicated that SSRIs could reach the ER freely.

Reapplication of the quaternary SSRI did not generate an increase in biosensor fluorescence signal over the existing baseline, which indicated that the quaternary compound still did not cross into the ER (Fig. 8, third row). Incubation of quaternary SSRIs with the PM-targeted biosensors (Fig. 8, fourth row) resembled the signals obtained with the 2.4 h incubation of the SSRIs with the PM-targeted biosensors (Fig. 8, second row).

When we attempted a 24 h pre-incubation with drug, we experienced a low ΔF/F_0_ that was confounded by high background (data not shown). We abandoned experiments with the 24 h pre-incubation.

### Membrane-impermeant SSRI derivatives are modestly weaker blockers

The membrane-impermeant quaternary SSRI derivatives provided an opportunity to test the hypothesis that the potency of SSRIs at SERT arises, in part, because they approach their binding site from the membrane phase. To compare the results with the time scale of our fluorescence experiments, we employed temporally resolved measurements on the transport-related current evoked by 5-HT (Mager et al., 1994), using an hSERT mutant that has unusually large transport-associated currents at low pH (Cao et al., 1997).

With membrane-bound hSERT in living cells, we found that N-N-dimethylfluoxetine blocks hSERT with an IC_50_ ∼11-fold higher than fluoxetine (Fig. 9A and C). With membrane-bound hSERT in living cells, we found that N-methylescitalopram blocks hSERT with an IC_50_ ∼ 6-fold greater than escitalopram (Fig 9. B and D). In more conventional experiments using [^3^H]serotonin flux, previous experiments found that a quaternary citalopram derivative blocks hSERT with a 10- fold higher IC_50_ than citalopram (Bismuth-Evenzal et al., 2010). These modest differences between the SSRIs and their quaternary derivatives do not strongly support the hypothesis that fluoxetine and escitalopram approach their binding site from the membrane (see Discussion).

### Intracellularly bioavailable fluoxetine and escitalopram equal the extracellular values but represent a small fraction of the total cellular drug due to lipid binding

To complement the iDrugSnFR experiments, we performed a series of measurements to measure both the ratio between the concentration of intracellular unbound (bioavailable) compound and that of the external solution (F_ic_), and the total cellular drug accumulation ratio (Kp). We employed cultured HEK293 cells, which provide a rough approximation (within 2-fold) to brain binding (Mateus et al., 2014).

We first describe the Kp data. Kp did not fully reach equilibrium for fluoxetine and escitalopram, with a peak at 30 mins and a subsequent decrease during the period leading up to 120 mins. At 30 and 120 min, K_p_ = 1590 ± 150, and 1170 ± 50 respectively (geometric mean ± SEM). The escitalopram K_p_ values were 12-fold smaller: at 30 and 120 min, 132 ± 12 and 67 ± 30 (geometric mean ± SEM) The values for K_p_ (Fig. 10A) are among the largest measured for any drug (Treyer et al., 2018).

**Figure 10.**
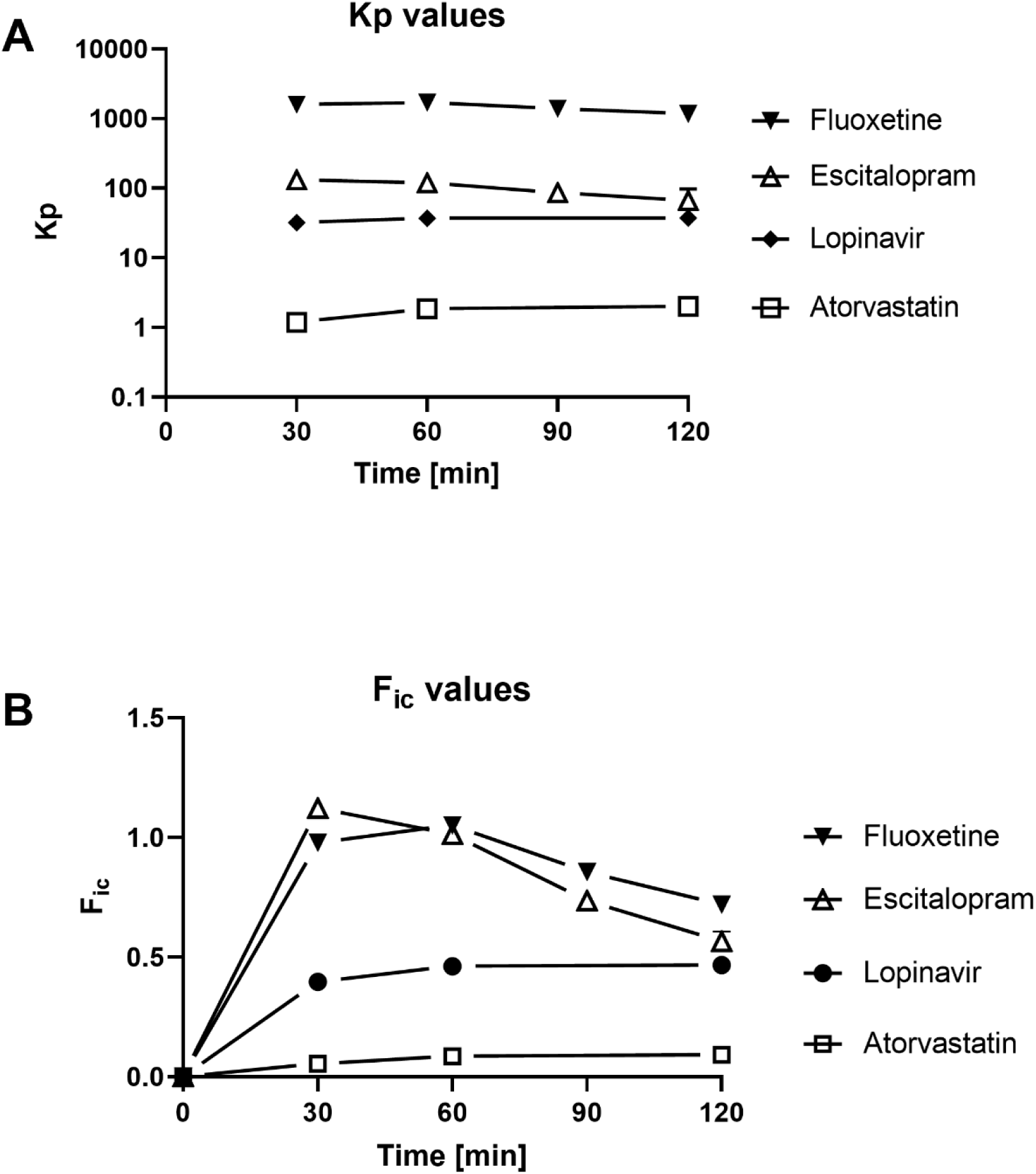
SSRIs are highly bioavailable intracellularly despite substantial membrane binding. **(A)** K_p_ values, measuring total cellular accumulation in living HEK293 cells at 30 to 120 min of incubation. **(B)** F_ic_ values, measuring the ratio between unbound intracellular (mostly cytoplasmic) concentration and the external solution. SEM values are shown where they exceed the size of data markers.

Recent experiments show that the intracellular unbound fraction of drug (f_u,cell_) is dominated by distribution into cellular membrane phospholipids (Treyer et al., 2018; Treyer et al., 2019). We determined f_u,cell_ for HEK293 cells with lipid membrane-coated beads using the approach developed by Treyer et al., 2019. The phosphatidylcholine coating substitutes approximately for measurements with phospholipid mixtures from individual cell types (Treyer et al., 2019), which were unavailable for these experiments. The experiments use a 12 min incubation, a time scale relevant to the iDrugSnFR experiments, The f_u,cell_ was 0.0006 for fluoxetine and 0.0085 for escitalopram. Thus, most of the intracellular drug is bound to lipids, and escitalopram binds less strongly than fluoxetine, consistent with the idea that iDrugSnFR kinetics for escitalopram were ∼ 10-fold faster than fluoxetine kinetics (Figs. 4 and 6) because diffusion of escitalopram through the membrane is buffered less by binding within the membrane.

The Kp and f_u,cell_ data then determined F_ic_ (Fig. 10B). F_ic_ ranged between 0.5 and 1.0 for both fluoxetine and escitalopram, after 60 min. This dataset was presumably dominated by cytoplasmically located drug, because the ER accounts for just ∼ 10% of total intracellular volume and other organelles represent even smaller volumes. The dataset thus agrees well with the similar ΔF/F_0_ measurements for the _cyto iDrugSnFRs vs free solution values. Because F_ic_ is proportional to K_p_, F_ic_ decreases, by ∼ 40-50%, between 60 min and 120 min. We have not systematically studied the origin of the decline, which was not observed for the two control drugs, atorvastatin and lopinavir.

Summarizing, chemical determination shows that applied fluoxetine or escitalopram enters the cell within 30 min. Of the intracellular SSRI, > 99% is bound to lipids and is therefore available for interaction with membrane proteins. Although < 1% of intracellular fluoxetine or escitalopram is unbound, this concentration roughly equals that of the external solution and is also available for interaction with SERT or other molecules.

## DISCUSSION

### Two SSRIs enter several cellular compartments

The present data establish that fluoxetine and escitalopram, two commonly used hSERT ligands, enter both the cytoplasm and the ER (the largest organelle) within at most a few min after the drugs appear outside a neuron (Fig. 4) or a HeLa cell (Fig. 6). The drugs leave with a similar time course after the extracellular [drug] is stepped to zero. That fluoxetine and escitalopram appear as unbound molecules at concentrations near the extracellular values is confirmed by chemical detection within HEK293 cells, a good model for intracellular pharmacokinetics of neurons (Mateus et al., 2013), albeit with less precise temporal resolution (Fig. 10).

At the same time the drugs are equilibrating with the cytoplasm and ER, the drugs are accumulating within the PM (Fig. 5, 7, and 10) and, presumably, the other membranes. We inferred the quantitative extent of accumulation within the membrane from iDrugSnFR waveforms. For fluoxetine sensed by iFluoxSnFR, our data for neurons and HeLa cells are consistent with concentration ratios of 180 (Fig. 5) and 18 (Fig. 6), respectively. The predicted logD_pH7.4_ value for fluoxetine corresponds to a concentration ratio of 67, but octanol and plasma membrane probably have different matrix properties. For pure phosphatidylcholine membranes on beads (this study), or for ITC measurement on pure 1,2-dierucoyl-*sn*-glycero-3-phosphocholine (Kapoor, 2019), the fluoxetine accumulation ratio is 30 – 300 times higher than we infer from iFluoxSnFR waveforms at the PM.

The data provide only slight support for the idea that both fluoxetine and escitalopram interact with SERT more strongly when approaching from the membrane than from an aqueous phase. The potency of membrane-impermeant quaternary derivatives as hSERT blockers is modestly (6- to 10-fold) less than the potency of the SSRIs themselves.

### Contributions of SSRI iDrugSnFRs to the iDrugSnFR paradigm

This study expands the iDrugSnFR family of sensors (Bera et al., 2019; Shivange et al., 2019; Muthusamy et al., 2022; Nichols et al., 2022) to include SSRIs. SSRI iDrugSnFRs are sensitive enough to allow experiments near the experimentally determined (or otherwise projected) concentration in human blood and CSF (Karson et al., 1992; Renshaw et al., 1992; Bolo et al., 2000; Paulzen et al., 2016).

Our previous applications of iDrugSnFRs have utilized such sensors to measure the free aqueous concentration of a drug. The present experiments show that at least one iDrugSnFR, iFluoxSnFR, can also detect fluoxetine in the membrane that anchors the iDrugSnFR. The iDrugSnFR experiments are well-suited to the apparent time scale of accumulation. This additional, useful feature presumably arises because the fluoxetine accumulates (our iDrugSnFR data are best fitted by a factor ∼181) in the membrane just a few Å from the binding site of iDrugSnFR. Similar accumulation in lipids also underlies the high volume of distribution (∼ 20 L/kg) that characterizes SSRIs in general. Other major antidepressant classes (tricyclics, serotonin-norepinephrine uptake inhibitors, and S-ketamine) have much lower volumes of distribution.

Some conventional measurements on SSRIs employ intracerebral microdialysis, with 10- 20 min sampling intervals (Fukushima et al., 2004; Bundgaard et al., 2007a; O’Brien et al., 2013) to examine 5-HT (or other neurotransmitter) levels (Cryan et al., 2004; Bundgaard et al., 2007b; Deltheil et al., 2009; Gardier, 2013). Neurotransmitter levels serve as an indirect indicator of SSRI concentration. Thus, imaging- or photometry-based examination of the local brain concentrations of local, free (unbound) antidepressant in real-time using iDrugSnFRs could provide valuable information on SSRI pharmacokinetics.

### Fluoxetine versus escitalopram

Our study exploited an important feature of a reduced cellular model: we performed experiments in parallel on two therapeutic agents thought to act similarly. The iDrugSnFRs themselves differ by only 8 amino acids near the PBP binding site and two near the PBP-cpGFP linkers (4 differences are shown in Fig. 1A, and full sequences are given in the Addgene deposits). The drugs themselves have rather similar logD_pH7.4_ and pKa values. We reasoned that any property governing SSRIs as a class would result in similar measurements for fluoxetine and escitalopram. We found two classes of shared properties: entry into the cytoplasm and ER, and accumulation in membranes.

Nonetheless, it is important to understand the different properties of fluoxetine and escitalopram. In medical practice, escitalopram produces fewer adverse events than fluoxetine, (Kennedy et al., 2009), and fluoxetine results in less frequent and less severe “antidepressant discontinuation syndrome” (Fava et al., 2015). All the present datasets are consistent with previous data that escitalopram accumulates in membranes or lipids roughly an order of magnitude less than fluoxetine (Wan et al., 2007; Lanevskij et al., 2011; Mateus et al., 2014; Kapoor et al., 2019). In the diffusion-bonding model, this difference explains how escitalopram enters and leaves the compartments we studied at least an order of magnitude faster than fluoxetine. The LogP and LogD_pH7.4_ for escitalopram are ∼ 0.5 less than for fluoxetine, perhaps corresponding to part of the difference in accumulation. The difference between fluoxetine and escitalopram is unlikely to arise from drug efflux pumps (Peters et al., 2009) but could arise from one or more other mechanisms, including kinetics of partitioning into lipid rafts (Senese and Rasenick, 2021), lateral diffusion within the plane of the membrane, and effects on membrane elasticity and curvature (Kapoor et al., 2019).

### Insights from quaternary SSRI derivatives

Use of impermeant quaternary blocking drugs is an accepted paradigm in ion channel and receptor pharmacology (Hille, 1977a, b; Shivange et al., 2019); and we performed analogous experiments with a neurotransmitter transporter. Some data suggest that fluoxetine stabilizes SERT in a conformation that exposes the binding site to the internal solution (Tavoulari et al., 2009); but atomic-scale structures suggest that bound escitalopram faces the external solution (Coleman et al., 2016). Neither of the two cited studies addresses the question of whether the SSRI approaches the binding site from the membrane or from the aqueous phase.

On the one hand, the modest decreases in affinity for the impermeant derivatives (6- to 10-fold) provide little support for the membrane approach mechanism. The amine of several SSRIs makes a cation-π interaction with Tyr95 and a hydrogen bond with Asp98 of SERT (Coleman and Gouaux, 2018). Quaternerizing the amine would alter the former interaction and eliminate the latter, possibly decreasing the affinity by the observed amounts. On the other hand, the impermeant derivatives will provide a convenient probe to distinguish effects of SERT blockade from intracellular effects of intracellular SSRI-SERT interactions, for at least 2.4 h (Fig. 8).

### Implications for the four possible mechanisms of SSRI action

Returning to the four non-exclusive mechanisms summarized in the Introduction: Mechanism (1), the “outside-in” mechanism, is not informed by our data. “Outside-in” processes operate via SSRI-driven changes in external 5-HT concentration, which we did not study.

Mechanism (2) - The hypothesis that SSRI levels during the therapeutic lag are governed by whole-animal or organ-level pharmacokinetic properties is not supported by our experiments— even if one assumes that myelin, with some 500 membranes in parallel, increases the wash-in and washout time constants for fluoxetine (300 s) by 500-fold. This would extend the times to 1.5 x 10^4^ s, or one day—enough to explain the classically measured disappearance of fluoxetine but still ∼ ten-fold less than the therapeutic lag. The faster kinetics for escitalopram further undercut the idea that lipid accumulation can explain the “therapeutic lag” for SSRIs.

However, use of SSRIs in premenstrual syndrome is apparently not associated with a “therapeutic lag” (Steinberg et al., 2012). The time course of antidepressant discontinuation syndrome is also relatively rapid compared with the classical therapeutic lag. Therefore, the purely pharmacokinetic hypothesis is being investigated further (Senese and Rasenick, 2021).

Mechanism (3) - The hypothesis that therapeutic effects occur at least partially because of SSRI-SERT interactions in cellular compartments other than the extracellular-facing surface of the PM, is consistent with our observations. Given the dimerization and quality control processes that transporters undergo in the ER, target engagement within the ER, including pharmacological chaperoning of nascent SERT, continues as a suspected therapeutic mechanism (Lester et al., 2012). That fluoxetine enters the ER may also explain how fluoxetine induces cytotoxic ER stress (Bowie et al., 2015). The vast SSRI accumulation within membranes and the decreased potency of membrane-excluded derivatives raises the possibility that SSRI-SERT engagement is enhanced because it occurs within the PM or an organellar membrane—a suggestion that broadens the meaning of the earlier phrase, “inside-out” (Lester et al., 2012). Presently available iDrugSnFRs cannot function in acidic organelles and therefore cannot enlighten the hypothesis of endosome-based SERT recycling (Riad et al., 2001; Riad et al., 2004).

Mechanism (4) - Additional pathways are consistent with our experiments to the extent that they involve SSRIs within membranes. Such pathways include interactions with TRKB (Casarotto et al., 2021), lipid rafts (Senese and Rasenick, 2021), or lipid-modifying enzymes (Kornhuber and Gulbins, 2021).

The hSERT ligands studied here have important continuing uses in medicine. These uses call for continuing investigations into the neuroscientific basis of their action(s).

## Acknowledgements

We thank Stefan Petrovic for his stewardship of the isothermal titration calorimeter in the Caltech Center for Molecular Medicine, the Gradinaru lab and Caltech CLOVER Center for help with viral vectors, and Andres Collazo and Giada Spigolon at the Caltech Biological Imaging Facility. We thank Zoe Beatty, Kallol Bera, Eve Fine, Shan Huang, Elaine Lin, Stephen Mayo, Lin Tian, Andrea Treyer, Elizabeth Unger, and Lu Wei for advice and guidance. We also thank Purnima Deshpande for her excellent lab management.

## Competing interests

The authors declare no financial or non-financial competing interests.

## Funding

California Tobacco-Related Disease Research Program (TRDRP) (27FT-0022), Aaron L. Nichols.

California Tobacco-Related Disease Research Program (TRDRP) (27IP-0057), Henry A. Lester.

California Tobacco-Related Disease Research Program (TRDRP) (T29IR0455), Dennis A. Dougherty.

NIH (GM-123582, MH120823), Henry A. Lester.

NIH (DA049140, GM7616), Anand K. Muthusamy.

Howard Hughes Medical Institute, Jonathan S. Marvin.

Howard Hughes Medical Institute, Loren L. Looger

Leiden University International Studies Fund (LISF L18020-1-45), Laura Luebbert.

Swedish Research Council, (01586), P. Artursson

European Union’s Horizon 2020 Research and Innovation Programme under the Marie Skłodowska-Curie grant agreement (956851), R. Hammar

## Notes

The authors declare no competing financial interests.

https://github.com/lesterha/lesterlab_caltech

### Competing Interest Statement

The authors have declared no competing interest.

## REFERENCES

Alberts B, Bray D, Lewis J, Morgan D, Raff M, Roberts K, Walter P, Wilson J, Hunt TW (2015) Molecular Biology of the Cell, 6th Edition. New York: Garland Science.

Armstrong D, Lester HA (1979) The kinetics of tubocarine action and restricted diffusion within the synaptic cleft. J Physiol 294:365–386.

Baldwin DS (2002) Escitalopram: efficacy and tolerability in the treatment of depression. Hosp Med 63:668–671.

Beasley CM, Jr., Nilsson ME, Koke SC, Gonzales JS (2000) Efficacy, adverse events, and treatment discontinuations in fluoxetine clinical studies of major depression: a meta-analysis of the 20-mg/day dose. J Clin Psychiatry 61:722–728.

Belmaker RH, Agam G (2008) Major depressive disorder. N Engl J Med 358:55–68.

Bera K, Kamajaya A, Shivange AV, Muthusamy AK, Nichols AL, Borden PM, Grant S, Jeon J, Lin E, Bishara I, Chin TM, Cohen BN, Kim CH, Unger EK, Tian L, Marvin JS, Looger LL, Lester HA (2019) Biosensors Show the Pharmacokinetics of S-Ketamine in the Endoplasmic Reticulum. Front Cell Neurosci 13:499.

Bismuth-Evenzal Y, Roz N, Gurwitz D, Rehavi M (2010) N-methyl-citalopram: A quaternary selective serotonin reuptake inhibitor. Biochem Pharmacol 80:1546–1552.

Bolo NR, Hode Y, Nedelec JF, Laine E, Wagner G, Macher JP (2000) Brain pharmacokinetics and tissue distribution in vivo of fluvoxamine and fluoxetine by fluorine magnetic resonance spectroscopy. Neuropsychopharmacology 23:428–438.

Borden P, Shivange AV, Marvin JS, Cichon J, Dan C, Podgorski K, Novak O, Tanimoto M, Lobas M, Khakh B, Dittman J, Gan W, Koyama M, Jayaraman V, Zhu J, Lester HA, Looger LL (2019) A genetically encoded fluorescent sensor for in vivo acetylcholine detection. bioRxiv.

Bowie M, Pilie P, Wulfkuhle J, Lem S, Hoffman A, Desai S, Petricoin E, Carter A, Ambrose A, Seewaldt V, Yu D, Ibarra Drendall C (2015) Fluoxetine induces cytotoxic endoplasmic reticulum stress and autophagy in triple negative breast cancer. World J Clin Oncol 6:299–311.

Bundgaard C, Jorgensen M, Larsen F (2007a) Pharmacokinetic modelling of blood-brain barrier transport of escitalopram in rats. Biopharm Drug Dispos 28:349–360.

Bundgaard C, Jorgensen M, Mork A (2007b) An integrated microdialysis rat model for multiple pharmacokinetic/pharmacodynamic investigations of serotonergic agents. J Pharmacol Toxicol Methods 55:214–223.

Burke WJ, Gergel I, Bose A (2002) Fixed-dose trial of the single isomer SSRI escitalopram in depressed outpatients. J Clin Psychiatry 63:331–336.

Cao Y, Mager S, Lester HA (1997) H^+^ permeation and pH regulation at a mammalian serotonin transporter. J Neurosci 17:2257–2266.

Casarotto PC et al. (2021) Antidepressant drugs act by directly binding to TRKB neurotrophin receptors. Cell 184:1299–1313 e1219.

Challis RC, Kumar SR, Chan KY, Challis C, Beadle K, Jang MJ, Kim HM, Rajendran PS, Tompkins JD, Shivkumar K, Deverman BE, Gradinaru V (2019) Publisher Correction: Systemic AAV vectors for widespread and targeted gene delivery in rodents. Nat Protoc 14:2597.

Clevenger SS, Malhotra D, Dang J, Vanle B, IsHak WW (2018) The role of selective serotonin reuptake inhibitors in preventing relapse of major depressive disorder. Ther Adv Psychopharmacol 8:49–58.

Coleman JA, Gouaux E (2018) Structural basis for recognition of diverse antidepressants by the human serotonin transporter. Nat Struct Mol Biol 25:170–175.

Coleman JA, Green EM, Gouaux E (2016) X-ray structures and mechanism of the human serotonin transporter. Nature.

Crank J (1975) The Mathematics of Diffusion, Second Edition. Oxford: Clarendon Press.

Cryan JF, O’Leary OF, Jin SH, Friedland JC, Ouyang M, Hirsch BR, Page ME, Dalvi A, Thomas SA, Lucki I (2004) Norepinephrine-deficient mice lack responses to antidepressant drugs, including selective serotonin reuptake inhibitors. Proc Natl Acad Sci U S A 101:8186–8191.

Deltheil T, Tanaka K, Reperant C, Hen R, David DJ, Gardier AM (2009) Synergistic neurochemical and behavioural effects of acute intrahippocampal injection of brain-derived neurotrophic factor and antidepressants in adult mice. Int J Neuropsychopharmacol 12:905–915.

Fava GA, Gatti A, Belaise C, Guidi J, Offidani E (2015) Withdrawal Symptoms after Selective Serotonin Reuptake Inhibitor Discontinuation: A Systematic Review. Psychother Psychosom 84:72–81.

Fukushima T, Naka-aki E, Guo X, Li F, Vankeirsbilck V, Baeyens WRG, Imaii K, Toshimasa T (2004) Determination of fluoxetine and norfluoxetine in rat brain microdialysis samples following intraperitoneal fluoxetine administration. Analytica Chimica Acta 522:99–104.

Gardier AM (2013) Antidepressant activity: contribution of brain microdialysis in knock-out mice to the understanding of BDNF/5-HT transporter/5-HT autoreceptor interactions. Front Pharmacol 4:98.

Gibson DG, Young L, Chuang RY, Venter JC, Hutchison CA, 3rd, Smith HO (2009) Enzymatic assembly of DNA molecules up to several hundred kilobases. Nat Methods 6:343–345.

Govind AP, Vallejo YF, Stolz JR, Yan JZ, Swanson GT, Green WN (2017) Selective and regulated trapping of nicotinic receptor weak base ligands and relevance to smoking cessation. Elife 6.

Guttler T, Madl T, Neumann P, Deichsel D, Corsini L, Monecke T, Ficner R, Sattler M, Gorlich D (2010) NES consensus redefined by structures of PKI-type and Rev-type nuclear export signals bound to CRM1. Nat Struct Mol Biol 17:1367–1376.

Hille B (1977a) The pH-dependent rate of action of local anesthetics on the node of Ranvier. J Gen Physiol 69:475–496.

Hille B (1977b) Local anesthetics: hydrophilic and hydrophobic pathways for the drug-receptor reaction. J Gen Physiol 69:497–515.

Kapoor R, Peyear TA, Koeppe RE, 2nd, Andersen OS (2019) Antidepressants are modifiers of lipid bilayer properties. J Gen Physiol 151:342–356.

Karson CN, Newton JE, Mohanakrishnan P, Sprigg J, Komoroski RA (1992) Fluoxetine and trifluoperazine in human brain: a 19F-nuclear magnetic resonance spectroscopy study. Psychiatry Res 45:95–104.

Kennedy SH, Andersen HF, Thase ME (2009) Escitalopram in the treatment of major depressive disorder: a meta-analysis. Curr Med Res Opin 25:161–175.

Kille S, Acevedo-Rocha CG, Parra LP, Zhang ZG, Opperman DJ, Reetz MT, Acevedo JP (2013) Reducing codon redundancy and screening effort of combinatorial protein libraries created by saturation mutagenesis. ACS Synth Biol 2:83–92.

Kornhuber J, Gulbins E (2021) New Molecular Targets for Antidepressant Drugs. Pharmaceuticals (Basel) 14.

Lalit V, Appaya PM, Hegde RP, Mital AK, Mittal S, Nagpal R, Palaniappun V, Ramsubramaniam C, Rao GP, Roy K, Trivedi JK, Vankar GK, Karan RS, Shah S, Patel RB (2004) Escitalopram Versus Citalopram and Sertraline: A Double-Blind Controlled, Multi-centric Trial in Indian Patients with Unipolar Major Depression. Indian J Psychiatry 46:333–341.

Lanevskij K, Dapkunas J, Juska L, Japertas P, Didziapetris R (2011) QSAR analysis of blood-brain distribution: the influence of plasma and brain tissue binding. J Pharm Sci 100:2147–2160.

Lee-Kelland R, Zehra S, Mappa P (2018) Fluoxetine overdose in a teenager resulting in serotonin syndrome, seizure and delayed onset rhabdomyolysis. BMJ Case Rep 2018.

Lepola UM, Loft H, Reines EH (2003) Escitalopram (10-20 mg/day) is effective and well tolerated in a placebo-controlled study in depression in primary care. Int Clin Psychopharmacol 18:211–217.

Lester HA, Miwa JM, Srinivasan R (2012) Psychiatric Drugs Bind to Classical Targets Within Early Exocytotic Pathways: Therapeutic Effects. Biol Psychiatry 72:905–915.

Lester HA, Lavis LD, Dougherty DA (2015) Ketamine Inside Neurons? Am J Psychiatry 172:1064–1066.

Lester HA, Xiao C, Srinivasan R, Son C, Miwa J, Pantoja R, Dougherty DA, Banghart MR, Goate AM, Wang JC (2009) Nicotine is a Selective Pharmacological Chaperone of Acetylcholine Receptor Number and Stoichiometry. Implications for Drug Discovery. AAPS Journal 11:167–177.

Loryan I, Friden M, Hammarlund-Udenaes M (2013) The brain slice method for studying drug distribution in the CNS. Fluids Barriers CNS 10:6.

Mager S, Min C, Henry DJ, Chavkin C, Hoffman BJ, Davidson N, Lester HA (1994) Conducting states of a mammalian serotonin transporter. Neuron 12:845–859.

Malhi GS, Mann JJ (2018) Depression. Lancet 392:2299–2312.

Marvin JS, Borghuis BG, Tian L, Cichon J, Harnett MT, Akerboom J, Gordus A, Renninger SL, Chen TW, Bargmann CI, Orger MB, Schreiter ER, Demb JB, Gan WB, Hires SA, Looger LL (2013) An optimized fluorescent probe for visualizing glutamate neurotransmission. Nat Methods 10:162–170.

Mateus A, Matsson P, Artursson P (2013) Rapid measurement of intracellular unbound drug concentrations. Mol Pharm 10:2467–2478.

Mateus A, Matsson P, Artursson P (2014) A high-throughput cell-based method to predict the unbound drug fraction in the brain. J Med Chem 57:3005–3010.

Muthusamy AK, Kim CH, Virgil SC, Knox HJ, Marvin JS, Nichols AL, Cohen BN, Dougherty DA, Looger LL, Lester HA (2022) Three Mutations Convert the Selectivity of a Protein Sensor from Nicotinic Agonists to S-Methadone for Use in Cells, Organelles, and Biofluids. J Am Chem Soc.

Nguyen PT, Lai JY, Lee AT, Kaiser JT, Rees DC (2018) Noncanonical role for the binding protein in substrate uptake by the MetNI methionine ATP Binding Cassette (ABC) transporter. Proc Natl Acad Sci U S A 115:E10596–e10604.

Nichols AL, Blumenfeld Z, Fan C, Luebbert L, Blom AEM, Cohen BN, Marvin JS, Borden PM, Kim C, Muthusamy AK, Shivange AV, Knox HJ, Campello HR, Wang JH, Dougherty DA, Looger LL, Gallagher T, Rees DC, Lester HA (2022) Fluorescence Activation Mechanism and Imaging of Drug Permeation with New Sensors for Smoking-Cessation Ligands. eLife 11:e74648.

Nierenberg AA, Farabaugh AH, Alpert JE, Gordon J, Worthington JJ, Rosenbaum JF, Fava M (2000) Timing of onset of antidepressant response with fluoxetine treatment. Am J Psychiatry 157:1423–1428.

O’Brien FE, O’Connor RM, Clarke G, Dinan TG, Griffin BT, Cryan JF (2013) P-glycoprotein inhibition increases the brain distribution and antidepressant-like activity of escitalopram in rodents. Neuropsychopharmacology 38:2209–2219.

Paulzen M, Lammertz SE, Grunder G, Veselinovic T, Hiemke C, Tauber SC (2016) Measuring citalopram in blood and cerebrospinal fluid: revealing a distribution pattern that differs from other antidepressants. Int Clin Psychopharmacol 31:119–126.

Peters EJ, Reus V, Hamilton SP (2009) The ABCB1 transporter gene and antidepressant response. F1000 Biol Rep 1:23.

Quan J, Tian J (2009) Circular polymerase extension cloning of complex gene libraries and pathways. PLoS One 4:e6441.

Rao N (2007) The clinical pharmacokinetics of escitalopram. Clin Pharmacokinet 46:281–290.

Renshaw PF, Guimaraes AR, Fava M, Rosenbaum JF, Pearlman JD, Flood JG, Puopolo PR, Clancy K, Gonzalez RG (1992) Accumulation of fluoxetine and norfluoxetine in human brain during therapeutic administration. Am J Psychiatry 149:1592–1594.

Riad M, Watkins KC, Doucet E, Hamon M, Descarries L (2001) Agonist-induced internalization of serotonin-1a receptors in the dorsal raphe nucleus (autoreceptors) but not hippocampus (heteroreceptors). J Neurosci 21:8378–8386.

Riad M, Zimmer L, Rbah L, Watkins KC, Hamon M, Descarries L (2004) Acute treatment with the antidepressant fluoxetine internalizes 5-HT1A autoreceptors and reduces the in vivo binding of the PET radioligand [18F]MPPF in the nucleus raphe dorsalis of rat. J Neurosci 24:5420–5426.

Salomon RM, Miller HL, Krystal JH, Heninger GR, Charney DS (1997) Lack of behavioral effects of monoamine depletion in healthy subjects. Biol Psychiatry 41:58–64.

Schapiro MB, Atack JR, Hanin I, May C, Haxby JV, Rapoport SI (1990) Lumbar cerebrospinal fluid choline in healthy aging and in Down’s syndrome. Arch Neurol 47:977–980.

Scheepers GH, Lycklama ANJA, Poolman B (2016) An updated structural classification of substrate-binding proteins. FEBS Lett 590:4393–4401.

Senese NB, Rasenick MM (2021) Antidepressants Produce Persistent Gα_s_-Associated Signaling Changes in Lipid Rafts after Drug Withdrawal. Mol Pharmacol 100:66–81.

Shivange AV, Borden PM, Muthusamy AK, Nichols AL, Bera K, Bao H, Bishara I, Jeon J, Mulcahy MJ, Cohen B, O’Riordan SL, Kim C, Dougherty DA, Chapman ER, Marvin JS, Looger LL, Lester HA (2019) Determining the pharmacokinetics of nicotinic drugs in the endoplasmic reticulum using biosensors. J Gen Physiol 151:738–757.

Smith D, Allerton C, Kalgutkar A, van de Waterbeemd H, Walker D (2012) Pharmacokinetics and Metabolism in Drug Design, 3rd Edition. Weinheim: Wiley.

Sogaard B, Mengel H, Rao N, Larsen F (2005) The pharmacokinetics of escitalopram after oral and intravenous administration of single and multiple doses to healthy subjects. J Clin Pharmacol 45:1400–1406.

Srinivasan R, Pantoja R, Moss FJ, Mackey EDW, Son C, Miwa J, Lester HA (2011) Nicotine upregulates α4β2 nicotinic receptors and ER exit sites via stoichiometry-dependent chaperoning. J Gen Physiol 137:59–79.

Steinberg EM, Cardoso GM, Martinez PE, Rubinow DR, Schmidt PJ (2012) Rapid response to fluoxetine in women with premenstrual dysphoric disorder. Depress Anxiety 29:531–540.

Tavoulari S, Forrest LR, Rudnick G (2009) Fluoxetine (Prozac) binding to serotonin transporter is modulated by chloride and conformational changes. J Neurosci 29:9635–9643.

Tischbirek CH, Wenzel EM, Zheng F, Huth T, Amato D, Trapp S, Denker A, Welzel O, Lueke K, Svetlitchny A, Rauh M, Deusser J, Schwab A, Rizzoli SO, Henkel AW, Muller CP, Alzheimer C, Kornhuber J, Groemer TW (2012) Use-dependent inhibition of synaptic transmission by the secretion of intravesicularly accumulated antipsychotic drugs. Neuron 74:830–844.

Treyer A, Walday S, Boriss H, Matsson P, Artursson P (2019) A Cell-Free Approach Based on Phospholipid Characterization for Determination of the Cell Specific Unbound Drug Fraction (fu,cell). Pharm Res 36:178.

Treyer A, Mateus A, Wisniewski JR, Boriss H, Matsson P, Artursson P (2018) Intracellular Drug Bioavailability: Effect of Neutral Lipids and Phospholipids. Mol Pharm 15:2224–2233.

Tucker KR, Block ER, Levitan ES (2015) Action potentials and amphetamine release antipsychotic drug from dopamine neuron synaptic VMAT vesicles. Proc Natl Acad Sci U S A 112:E4485–4494.

Unger EK et al. (2020) Directed Evolution of a Selective and Sensitive Serotonin Sensor via Machine Learning. Cell 183:1986–2002.e1926.

Vargas HM, Jenden DJ (1996) Elevation of cerebrospinal fluid choline levels by nicotinamide involves the enzymatic formation of N1-methylnicotinamide in brain tissue. Life Sci 58:1995–2002.

Wan H, Rehngren M, Giordanetto F, Bergström F, Tunek A (2007) High-throughput screening of drug-brain tissue binding and in silico prediction for assessment of central nervous system drug delivery. J Med Chem 50:4606–4615.

Wong DT, Bymaster FP, Engleman EA (1995) Prozac (fluoxetine, Lilly 110140), the first selective serotonin uptake inhibitor and an antidepressant drug: twenty years since its first publication. Life Sci 57:411–441.

Wong DT, Perry KW, Bymaster FP (2005) Case history: the discovery of fluoxetine hydrochloride (Prozac). Nat Rev Drug Discov 4:764–774.

Wong DT, Horng JS, Bymaster FP, Hauser KL, Molloy BB (1974) A selective inhibitor of serotonin uptake: Lilly 110140, 3-(p-trifluoromethylphenoxy)-N-methyl-3-phenylpropylamine. Life Sci 15:471–479.

Yu H, Dickson EJ, Jung SR, Koh DS, Hille B (2016) High membrane permeability for melatonin. J Gen Physiol 147:63–76.

Zeisel SH, Epstein MF, Wurtman RJ (1980) Elevated choline concentration in neonatal plasma. Life Sci 26:1827–1831.

